# A quantitative proteomics dataset for assessment and prediction of low dose X-ray radiation exposure in mice

**DOI:** 10.64898/2026.05.18.725951

**Authors:** Alex Zelter, Michael Riffle, Gennifer E. Merrihew, Batool Mutawe, Nicholas Shulman, Justin A. Sanders, William S. Noble, Danielle P. Johnson Erickson, Alec Morimoto, Benjamin A. Shaver, Taylor N. Steins, Ning Cao, Eric C. Ford, Paul A. Rudnick, Daniel Chelsky, Kenneth H. Wan, Jamie L. Inman, Hang Chang, Antoine M. Snijders, Jian-Hua Mao, Susan E. Celniker, Jared De Chant, Lieselotte Obst-Huebl, Kei Nakamura, Christine C. Wu, Michael J. MacCoss

**Affiliations:** Department of Genome Sciences, University of Washington, Seattle, WA, USA; Department of Radiation Oncology, University of Washington School of Medicine, Seattle, WA, USA; Spectragen Informatics LLC, Bainbridge Island, WA, USA; Biological Systems and Engineering Division, Lawrence Berkeley National Laboratory, 1 Cyclotron Rd, Berkeley, CA, USA; Biosciences and Biotechnology Division, Lawrence Livermore National Laboratory, 7000 East Avenue, Livermore, CA, USA; Accelerator Technology & Applied Physics Division, Lawrence Berkeley National Laboratory, 1 Cyclotron Rd, Berkeley, CA, USA

## Abstract

Ionizing radiation induces molecular responses that may be used to estimate exposure when physical dosimeters are unavailable. Here we present two large-scale proteomics datasets generated from mouse dorsal skin punch samples collected following controlled X-ray exposures spanning multiple doses, dose rates, and post-exposure time points. Experiment 1 comprised 96 samples (including 16 reference samples) collected 6 days after exposure to 0-75 cGy delivered at either 30 or 300 cGy/min. Experiment 2 comprised 936 samples (including 236 reference samples) exposed to 0-100 cGy at either 3 or 28 cGy/min dose rates and harvested between 7 and 150 days post-exposure. All samples were processed using a standardized workflow involving automated bead-based digestion and data-independent acquisition mass spectrometry. The datasets include multiple pooled reference sample types, process controls, and system suitability standards ensuring high quality data. All data presented are available via ProteomeXchange at several levels of processing, from raw files through normalized peptide- and protein-level abundance matrices suitable for biomarker discovery and machine learning applications. This dataset will facilitate generation of new insights into the biological changes and molecular signatures resulting from X-ray exposure in mice and may also help inform future studies in humans.

## Background & Summary

Accurate assessment of ionizing radiation exposure is essential for medical management, occupational monitoring, and response to radiological or nuclear incidents. When physical dosimeters are unavailable, biological dosimetry methods must be used to estimate radiation exposure by measuring biological responses that correlate with absorbed dose. Cytogenetic assays, particularly analysis of dicentric chromosomes in peripheral blood lymphocytes, remain the gold standard in biodosimetry approaches.^1^ However, these assays are labor-intensive and require specialized expertise, limiting their scalability in large-scale exposure scenarios.^2^ Such limitations provide motivation for the development of molecular biomarker based approaches capable of detecting and quantifying radiation exposure using accessible biological specimens.

Ionizing radiation induces a wide range of molecular changes within exposed tissues, including DNA damage, oxidative stress, and alterations in protein expression and post-translational modifications.^3–6^ Mass spectrometry (MS) based proteomic measurements provide a powerful means of capturing these molecular responses, as changes in protein abundance and modification state can reflect downstream biological responses to radiation exposure. Recent advances in data-independent acquisition MS (DIA-MS) now enable reproducible quantification of thousands of proteins across large sample cohorts, providing the opportunity to identify radiation-associated molecular signatures.^7,8^

Skin represents an attractive tissue for biodosimetry applications because it can be easily sampled using minimally invasive procedures and is directly exposed during many radiation exposure scenarios. Proteomic profiling of skin samples offers the potential to identify molecular signatures that correlate with radiation dose, dose rate and time after exposure. However, development of robust biomarker models requires large datasets generated under controlled experimental conditions, including well-defined radiation dose series and rigorous evaluation of experimental reproducibility.

Here we present two large-scale proteomics datasets generated from mouse dorsal skin punch samples collected following controlled X-ray radiation exposures across a range of doses, dose rates, and post-exposure time points (Figure 1). These datasets were produced as part of efforts to discover and evaluate molecular biomarkers capable of detecting and quantifying low-dose radiation exposure.

**Figure 1.**
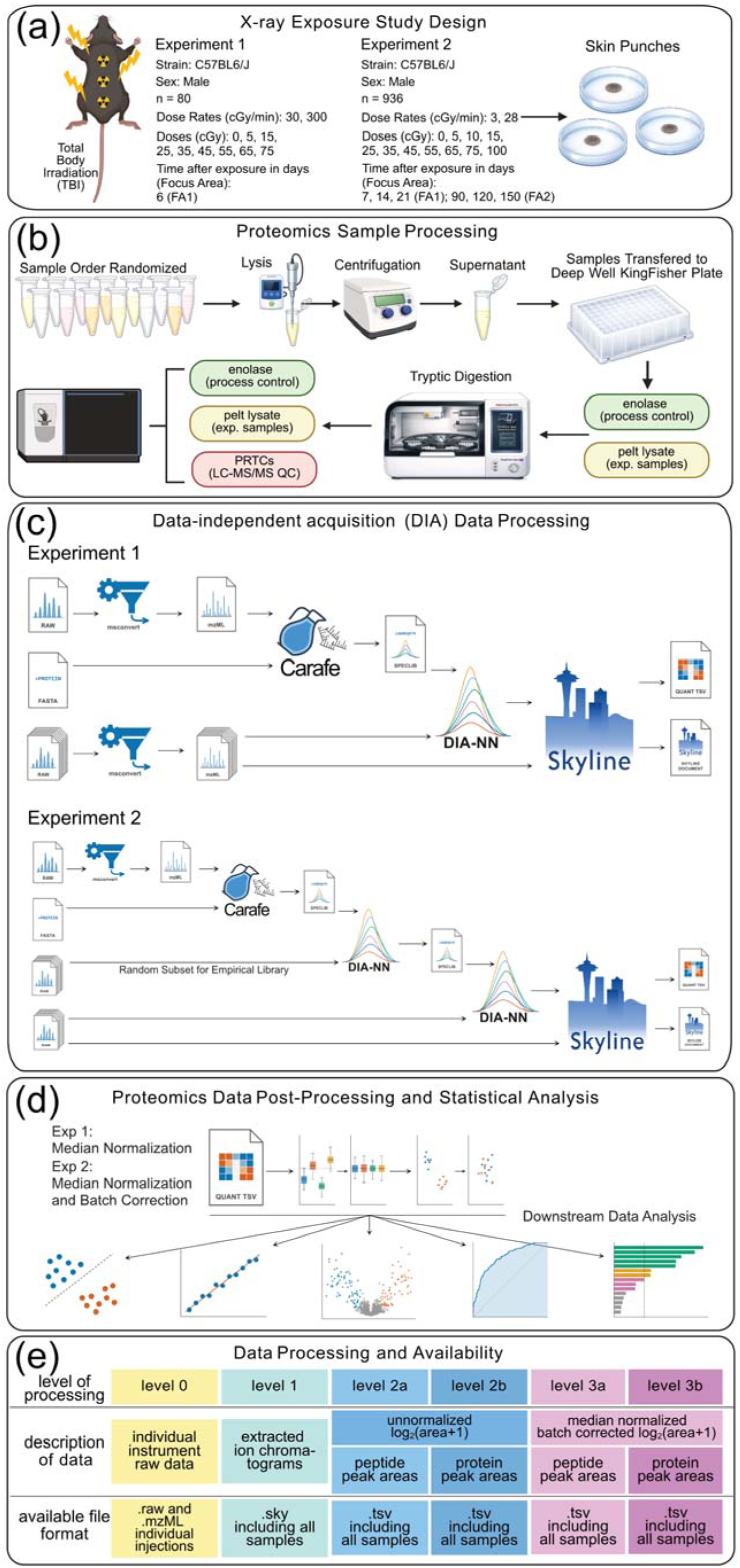
Overview of the X-ray exposure study design and associated proteomics workflow. (a) Mouse pelt tissue was collected from animals following X-ray exposure at doses ranging from 5 to 100 cGy and at time points from 6 to 150 days post-exposure. (b) Samples were randomized, processed, and analyzed in batches of up to 96 samples. Yeast enolase was added to each protein lysate as a process control prior to digestion. Following digestion, a Pierce PRTC synthetic peptide mixture was added as a system suitability control. (c) Individual samples were analyzed by single injections using wide-window DIA for quantitative analysis. Spectral libraries were generated using Carafe and peptide detection and scoring were performed using DIA-NN, with peak boundary refinement and integration carried out in Skyline. (d) Peptide and protein peak area data were evaluated, normalized, batch corrected and processed using in house code available on GitHub. (e) The proteomics data is publicly available on the Panorama web server at 5 different levels of processing.

Two complementary experiments are included. Experiment 1 is a smaller study comprising 80 samples (96 including the reference samples described below), in which mice were exposed to X-ray doses ranging from 0 to 75 cGy at dose rates of either 30 or 300 cGy/min. Skin punch samples were collected 6 days post-exposure and analyzed by DIA-MS to characterize dose-dependent proteomic responses.

Experiment 2 consists of a much larger experiment comprising 700 samples (936 including the reference samples described below), in which mice were exposed to doses ranging from 0 to 100 cGy at dose rates of 3 cGy/min (low dose rate, LDR) or 28 cGy/min (high dose rate, HDR) and harvested between 7 and 150 days post-exposure. The 700 samples were assigned a “Focus Area”, which partitioned samples by post-irradiation timepoint: FA1 covers early times (6–21 days) and FA2 covers late times (90–150 days). QC reference samples were not assigned to a Focus Area.

To enable assessment of experimental reproducibility and facilitate downstream normalization and batch correction, both experiments incorporated two independent pooled reference sample types, with one of each included in every row of each 96-well plate throughout sample preparation and data acquisition. Experiment 2 additionally incorporated a third reference type consisting of all 80 samples from Experiment 1. These samples were re-digested and reanalyzed within Experiment 2 and randomly interspersed across all 10 plates. They were included as positive controls to enable evaluation of the performance of the larger Experiment 2 dataset in capturing radiation-associated proteomic phenotypes.

The complete Experiment 2 dataset comprises 936 samples generated across ten 96-well plates using a standardized sample preparation workflow and were analyzed by DIA-MS^9–11^. DIA enables untargeted, quantitative characterization of the proteome in complex biological samples. In DIA, extracted precursor-to-product ion chromatograms (XICs) can be generated for all detectable peptides across the sampled *m/z* range, enabling quantitative analysis without the need for pre-specified protein or peptide targets. We performed DIA analysis on all samples, enabling the detection and differential analysis of all peptides present within the acquired *m/z* range.

The design of the experiments presented here enables the discovery and evaluation of proteomic biomarkers associated with radiation exposure while also providing extensive internal controls for assessing analytical reproducibility and normalization strategies across large proteomics datasets.

Together, these datasets provide a valuable resource for investigating molecular responses to low-dose radiation exposure, developing predictive models for radiation biodosimetry, and evaluating computational approaches for normalization and batch correction in large-scale proteomics studies. The scale of the dataset, combined with its controlled experimental design and extensive reference samples, makes it well suited for machine learning applications and biomarker discovery efforts aimed at improving radiation exposure assessment.

## Methods

### Experimental Design

For Experiment 1, animals were caged in groups of 4 and were randomly assigned by cage to experimental groups. For Experiment 2, group sizes were determined a priori based on power calculations and animals were randomly assigned to experimental groups by cage. Animals were monitored daily for general health and signs of distress throughout the study. Any exclusions, human endpoints, and criteria for removal from analysis were predefined.

### Mouse Pelt Sample Production

#### Experiment 1

##### Animals

All animal procedures were conducted in accordance with institutional guidelines and were approved by the Institutional Animal Care and Use Committee (IACUC) at the University of Washington under protocol number 4528-01. Male C57BL/6 11-week-old mice were sourced from The Jackson Laboratory and housed under standard conditions with food and water *ad libitum*. Animals were acclimated for 9 days prior to radiation exposure.

##### X-ray exposure

Animals received a single total body dose of 5, 15, 25, 35, 45, 55, 65, or 75 cGy using a Small Animal Radiation Research Platform (SARRP) Xray source (Xstrahl, Inc) under anesthesia with an intraperitoneal injection of ketamine (100-125 mg/kg) and xylazine (10-12.5 mg/kg) in sterile saline. Dose rates were calibrated based upon the procedures described in American Association of Physicist in Medicine (AAPM) Task Group Report 61 (TG-61)^12,13^ with regards to the following conditions: X-ray tube potential was 220 kV and the half value layer (HVL) was 0.67 mm copper (Cu) with a nominal field size of 13x17 cm at the treatment plane. Doses were measured at 0.5 cm depth in solid water phantom using a Preston-Tonks-Wallace (PTW) model N30013 Farmer-type ionization chamber and Standard Imaging, Inc, model SUPERMAX electrometer. The chamber and electrometer underwent calibration at the University of Wisconsin Medical Radiation Research Center Accredited Dosimetry Calibration Laboratory (UWMRRC ADCL) using a beam quality of UW200-M and UW150-M. Two different dose rates were used: 300 cGy/min (tube current 10.7 mA, source-to-surface distance 33 cm) and 30 cGy/min (4.6 mA, SSD 76 cm). Control animals were sham irradiated under anesthesia.

##### Sample collection

Animals were euthanized by cervical dislocation under isoflurane anesthesia 6 days post-irradiation. A minimum of n = 4 mice were collected per exposure. Immediately following euthanasia, mice were shaved and pelts were dissected using a dorsal midline incision. 5-mm skin punches were collected from the dissected pelts using disposable biopsy punches (Integra Miltex, catalog #33-35), transferred into sterile 1.5 mL microcentrifuge tubes, frozen on dry ice, and stored at −80°C.

#### Experiment 2

##### Animals

All animal procedures were conducted in accordance with institutional guidelines and were approved by the Institutional Animal Welfare and Research Committee at Lawrence Berkeley National Laboratory (LBNL) under protocol number 270036. All studies were performed in accordance with the Guide for the Care and Use of Laboratory Animals. Male C57BL/6 (Jackson Laboratory, strain 000664) mice aged 6-8 weeks were purchased from The Jackson Laboratory. Animals were maintained under specific pathogen-free conditions under controlled environment at 21.9 ± 2°C and 54.3 ± 10% relative humidity on a 12-hour light/12-hour dark cycle. Mice had *ad libitum* access to 5053 PicoLab Rodent Diet 20 and water. Animals were acclimated for at least two weeks prior to radiation exposure.

##### X-ray exposure

At 9-12 weeks of age, mice were anesthetized with an intraperitoneal injection of ketamine (100-125 mg/kg) and xylazine (10-12.5 mg/kg) in sterile saline. Anesthetized mice were exposed to whole-body X-ray irradiation using a Precision X-ray XRAD320 system (Precision X-Ray, Inc., North Branford, CT) operated at 300 kVp. The irradiation setup included a collimator, motorized table, and a rotation stage (4 RPM) to ensure uniform dose delivery. The instrument was equipped with tri-metal (1.5mm Al, 0.25mm Cu, 0.75mm Sn) hardening filter. Animals were positioned 72 cm from the X-ray source dose rates were controlled by adjusting the current (1 mA = 0.03 Gy/min; 10 mA = 0.282 Gy/min). Dosimetry was performed using a RadCal ion chamber (Radcal 10X6-0.18). The transverse dose profile was measured using EBT3 radiochromic films with the collimator fully open and showed that within the radius of 110 mm, the dose profile was uniform within 4% (standard deviation) across the field.

Dose harmonization, between UW and LBNL, was performed using EBT3 radiochromic film and showed cross-site dose agreement within 12% of expected values, with an average deviation of approximately 6% below the nominal dose (data not shown).

##### Sample collection

Post radiation exposure, animals were randomly assigned to different groups for sample collection at 7, 14, 21, 90, 120, and 150 days. Groups were “focus area 1” (FA1), which covered early times (6–21 days) and FA2 which covered late times (90–150 days). Blood was collected by retroorbital bleeding under anesthesia, after which mice were euthanized. Following euthanasia, mice were dissected and the skin was separated from the body cavity. Up to eight 5-mm pelt biopsies were collected along the dorsal surface of each mouse. Individual biopsies were placed into 1.5mL Protein LoBind (Eppendorf, product number 022431081) and flash frozen in liquid nitrogen. Samples were stored at −80°C until processing for shipment. For shipment, samples were organized on dry ice, packed in shipping boxes with dry ice, and sent overnight to the University of Washington.

##### Humane endpoints and monitoring

Animals were monitored daily for signs of pain or distress. Humane endpoints included weight loss of 15% or poor body condition, defined as a body condition score 2 or lower.

### Mouse Pelt Lysate Production

Pelt punches were processed in single plate batches. Each plate consisted of up to 96 samples. Experiment 1 was a single plate. Experiment 2 consisted of 10 plates. All samples were randomized before processing for proteomics analysis, and the plate number and sample order were recorded for each sample in the metadata for each experiment as described in the Technical Validation, Experimental Design and Internal Controls section below.

Individual pelt punches were transferred to LoBind Eppendorf tubes, followed by addition of 250 µL of lysis buffer (100 mM Tris, pH 8.5, 2% SDS) supplemented with HALT Protease and Phosphatase Inhibitor Single-Use Cocktail (Thermo Scientific, Cat. No. PI78442). Samples were sonicated using a Misonix S-400 equipped with a microtip at 34% amplitude (6.75 kHz) for 10 seconds, followed by four cycles at 40% amplitude (8 kHz) for 30 seconds with 30-second rest intervals on ice, resulting in a total sonication time of 2 to 4 minutes. The sonicator probe was cleaned with 70% ethanol between samples. Once all samples were sonicated, the resulting lysates were centrifuged at 1,000 × g for 10 minutes, after which the supernatant was transferred to a fresh LoBind Eppendorf tube.

For Experiment 1, two pooled reference lysates were also generated. The first was a batch reference consisting of a pooled lysate generated from all 80 experimental lysates included in Experiment 1, which is labeled as “Batch Ref” in the metadata, (see Supplementary Table S1; Supplementary_Table_S1.xlsx for metadata headings and https://panoramaweb.org/teirex-2a-ldxr-mouse-pelt.url for complete metadata). The second, labeled “batch QC” in the metadata, was generated from a previous experiment not otherwise described in this study and that consisted of pooled lysates from 48 male BALB/c mice (14 weeks old) exposed to 0, 0.75, or 2 Gy low-dose neutron radiation and harvested 6 days after exposure.

For Experiment 2, two pooled reference lysates were similarly generated. The inter-batch reference, labeled “UW IB Reference” in the metadata, (see Supplementary Table S2; Supplementary_Table_S2.xlsx for metadata headings and https://panoramaweb.org/teirex-2a-ldxr-mouse-pelt.url for complete metadata), consisted of pooled lysates from all experimental samples present on plates 1 and 2 of Experiment 2. The inter-experiment reference (labeled “UW IE Reference” in the metadata) consisted of pooled lysates from a separate prior experiment not otherwise described in this study and included samples from 32 male C57BL/6J mice (22 weeks old) exposed to 0, 4, or 18 Gy proton or X-ray radiation and harvested 6 or 25 days after exposure. Experiment 2 additionally incorporated a third “internal control” reference type consisting of all 80 samples from Experiment 1. These lysates were re-digested and reanalyzed within Experiment 2 and are labeled “UW IC Reference” in the metadata.

For both Experiments 1 and 2, each sample lysate was vortexed, and 50 µL of lysate was transferred to the appropriate well of a KingFisher 96-well plate (Thermo Fisher Scientific, cat. no. 95040450). In addition to experimental samples, reference lysates were aliquoted into wells throughout the plates as described above. Once an entire KingFisher plate was complete, the plate was sealed using a SureSTART WebSeal 96-Well Plate Sealing Mat (Thermo Fisher Scientific, cat no. 60180-M185) followed by a Polystyrene Universal Microplate Lid (Corning, cat no. 3099), which was further sealed with Parafilm Laboratory Film. This plate was stored at −80°C prior to further workup for MS, which was performed within 5 days of initial pipetting.

### Sample Preparation

Previously prepared KingFisher plates containing 50 µL of mouse pelt lysate per well were removed from −80°C and thawed at 37°C in an Eppendorf ThermoMixer (approximately 15 minutes). To each well, 40 µL of a solution containing yeast enolase (Sigma, cat no. E6126; added as a process control)^14^ and tris(2-carboxyethyl)phosphine (TCEP; Invitrogen, Bond-Breaker TCEP Solution, Neutral pH, cat no. 77720) in 100 mM Tris buffer (pH 8.5, 2% SDS) was added, resulting in 800 ng enolase and 10 mM TCEP per well.

Plates were covered with a lid (Corning, cat no. 3099), mixed at 1,200 rpm for 30 seconds, and reduced at 37°C for 1 hour in an Eppendorf ThermoMixer with shaking at 650 rpm. Iodoacetamide (10 µL of 150 mM; Sigma, cat no. I1149) was then added to each well for a final concentration of 15 mM, and samples were alkylated at room temperature in the dark for 30 minutes. The reaction was quenched by addition of 10 µL of 150 mM DTT to each well.

MagResyn Hydroxyl particles (12.5 µL per well; ReSyn Biosciences, cat no. MR-HYX002) were added, and proteins were aggregated onto the beads by addition of 878 µL of a 1:1 ethyl acetate/ethanol mixture, yielding final solvent concentrations of approximately 44% ethyl acetate and 44% ethanol and a final volume of 1 mL per well. Beads were washed using five KingFisher deep-well plates (1 mL per well; three washes with 95% acetonitrile followed by two washes with 70% ethanol), with each wash step performed for 2.5 minutes using a KingFisher Apex (Thermo Fisher Scientific).

Protein-containing beads were transferred into 150 µL of 50 mM Tris (pH 8.5) containing 2.5 µg trypsin (Pierce, cat no. PI90058) per well, corresponding to an approximate 1:20 enzyme-to-substrate ratio assuming ∼50 µg protein per well. Digestion was performed at 47 °C for 1 hour in the KingFisher.

Following digestion, 100 µL of each sample was transferred to a Thermo Scientific SureSTART autosampler plate (cat no. 60180-P207B) containing 7 µL of 10% trifluoroacetic acid (TFA) per well and mixed by pipetting. A 50 µL aliquot of acidified peptides was then transferred to a new autosampler plate containing 5 µL of 500 fmol/µL Pierce Peptide Retention Time Calibration Mixture (PRTC; Thermo Fisher Scientific, cat no. PI-88321) in 0.1% TFA, added as a process control.^14^ After mixing, 20 µL of each sample was transferred to a final autosampler plate. For multi-plate experiments one plate was digested per day.

For Experiment 1, samples were distributed across two final autosampler plates, generating two half plates to reduce the amount of time individual samples remained in the autosampler at 7 °C prior to analysis. This was necessary because Experiment 1 was acquired on a slower LC–MS/MS platform (a Thermo Fisher Scientific Orbitrap Eclipse Tribrid) with longer analysis times per sample. In contrast, Experiment 2 was acquired on a faster instrument (a Thermo Fisher Scientific Astral) with shorter gradients, allowing full autosampler plates to be analyzed while maintaining similar sample residence times in the autosampler.

All buffers and solvents for the entire multi-plate Experiment 2 were prepared beforehand and were kept constant across the entire experiment. A single batch (one lot number) of trypsin and PRTCs were used across the entire experiment. Likewise – plastics were restricted to constant lot numbers across the entire experiment to minimize potential batch effects.

### Mass Spectrometry

Prepared samples were analyzed by DIA-MS. Each LC-MS/MS sample injection for Experiment 1 consisted of 6 µL containing a target amount (see “Mouse Pelt Lysate Production” above) of approximately 2 µg total mouse pelt protein plus 32 ng yeast enolase and 300 fmol of heavy labelled Pierce PRTC Mixture. Experiment 2 samples were prepared identically but analysis was performed on 3 µL sample injections and thus contained approximately 1 µg total mouse pelt protein plus 16 ng yeast enolase and 150 fmol of heavy labelled Pierce PRTC Mixture. Enolase and PRTCs were added to all samples as process controls.^14^ Reversed phase liquid chromatography was performed with a Thermo Vanquish Neo UHPLC system using a “trap and elute” configuration. A Thermo Scientific PepMap Neo Trap Cartridge (Cat. No. 174500) was used in all experiments. A 15 cm Bruker PepSep C18 column (15 cm × 150 µm, 1.9 µm; Cat. No. 1893471) was used for Experiment 1, while an 8 cm Bruker PepSep C18 column (8 cm × 150 µm, 1.5 µm; Cat. No. 1893470) was used for Experiment 2. Separate System Suitability LC-MS/MS runs^14^ were performed by injecting 6 µL (Experiment 1) or 3 µL (Experiment 2) of a peptide mix containing 50 fmol/µL PRTCs and 200 fmol/µL BSA in 0.1% TFA.

The HPLC column output was connected to a 5 cm × 20 µm ID Sharp Singularity Fossil Ion Tech tapered tip enclosed in a custom made microspray source heated to 45°C. For Experiment 1, peptides were eluted from the column at 0.8 µL/min using the following acetonitrile gradient: (1) 0-0.7 mins; 3-5% B; flow 1.3 µL/min; (2) 0.7-1 mins; 5-5.5% B; flow 1.3 µL/min; (3) 1-57 mins; 5.5-40% B; flow 0.8 µL/min; (4) 57-57.5 mins; 40-55% B; flow 1.3 µL/min; (5) 57.5-58 mins; 55-99% B; flow 1.3 µL/min; (6) 58-60 mins; 99% B; flow 1.3 µL/min.

For Experiment 2, peptides were eluted at 1 µL/min using the following acetonitrile gradient: (1) 0-0.7 mins; 3-4% B; flow 1.3 µL/min; (2) 0.7-1 mins; 4-4.5% B; flow 1.3 µL/min; (3) 1-22 mins; 4.5-38% B; flow 1 µL/min; (4) 22-22.5 mins; 38-50% B; flow 1 µL/min; (5) 22.5-23 mins; 50-99% B; flow 1.3 µL/min; (6) 23-24 mins; 99% B; flow 1.3 µL/min. For both experiments buffer A was 0.1% formic acid in water and buffer B was 0.1% formic acid, 80% acetonitrile and 20% water.

For Experiment 1, DIA data were collected on individual samples using a Thermo Fisher Scientific Orbitrap Eclipse Tribrid mass spectrometer. DIA was performed using a method with 12 *m/z* staggered precursor isolation windows.^15,16^ The Orbitrap resolving power for MS1 and MS/MS was 30,000 at *m/z* 200. Each cycle consisted of 1 MS scan followed by 50 MS/MS scans. MS scan range was *m/z* 395–1,005. Automatic gain control was set to a normalized value of 100% for MS scans and 800% for MS/MS scans. A normalized collision energy of 27 assuming charge state 3 was used for all MS/MS scans. Cycle 1 spanned 406.4347 *m/z* through 994.7021 *m/z* in 12 *m/z* isolation windows. Cycle 2 spanned 400.4319 *m/z* through 1000.7048 *m/z* in 12 *m/z* windows. All spectra were collected in centroid mode.

For Experiment 2, DIA data on individual samples were acquired using a Thermo Fisher Scientific Astral mass spectrometer. DIA was performed using 4 *m/z* non-staggered precursor isolation windows. The Orbitrap resolving power for MS1 was 240,000 at *m/z* 200. Each cycle consisted of 1 MS scan followed by 124 MS/MS scans. MS scan range was *m/z* 375–985. Automatic gain control was set to standard for MS scans and to a normalized value of 200% for MS/MS scans. A normalized collision energy of 27 assuming charge state 2 was used for all MS/MS scans. MS/MS isolation windows 400.4319-404.4337 through 896.6575-900.6593 *m/z*. All spectra were collected in centroid mode.

### Mass Spectrometry Data Conversion and Searching

For Experiment 1, acquired raw files were processed by an automated Nextflow^17^ workflow (https://github.com/mriffle/nf-skyline-dia-ms, git revision: c47c8346). The complete pipeline.config file is available on Panorama (https://panoramaweb.org/teirex-2a-ldxr-mouse-pelt.url), which results in the following tasks being performed (see Figure 1c): Raw files were demultiplexed^15,16^ and converted to mzML format in the workflow using ProteoWizard’s msConvert^18^ version 3.0.25168 using the following arguments: ‘**`--mzML --zlib --ignoreUnknownInstrumentError --filter “peakPicking true 1-” --64 --filter “demultiplex optimization=overlap_only” –simAsSpectra`**’. A fine-tuned spectral library was generated by Carafe^19,20^ version 2.0.0 using a FASTA file containing the UniProt reviewed mouse proteome (UP000000589), downloaded December 2024, and 2024_09_22_TRX_Phase2_TrainingPlate_Ecl_neo_TRX_LDXR_11_B11_32.mzML as input. The raw files were searched using the generated spectral library with DIA-NN^21^ version 2.2.0. DIA-NN results were imported into Skyline^22,23^ (Skyline-daily (64-bit: automated build) 25.1.1.168) using the command line, where the results were analyzed using Skyline’s native the “Impute boundaries for missing peaks” function (see details in the section below). Protein grouping was performed by Skyline (as described here: https://skyline.ms/wiki/home/software/Skyline/page.view?name=Skyline%20Protein%20Association%2022.1) using logic based on the IDPicker algorithm whereby proteins that match the same set of peptides are merged into indistinguishable protein groups.^24,25^ The precursor and protein quantities were then exported from Skyline for downstream analysis.

For Experiment 2, acquired raw files from 10 plates, each containing 96-wells, were processed as a single job using the same automated Nextflow workflow as for Experiment 1. The complete pipeline.config file for Experiment 2 data processing is available on Panorama, which results in a similar set of steps being performed as described above for Experiment 1. Differences from the Experiment 1 analysis are as follows (see Figure 1c): A fine-tuned spectral library was generated by Carafe using the same FASTA but with the Experiment 2 wide window file TRX_Phase2_Pelt-P01_Ast_Neo_TRX-TE-MSP-2002_007.mzML as input. This single mzML file was generated using ProteoWizard’s msConvert version 3.0.25168 using the following arguments: ‘**`--mzML --zlib --ignoreUnknownInstrumentError --filter “peakPicking true 1-” --64 –simAsSpectra`**’. An empirical spectral library was made from the Carafe library by running DIA-NN on 4 random raw files from each plate. All raw files were then searched with DIA-NN using this empirical spectral library. DIA-NN results were imported into Skyline where the results were processed and exported on a plate-by-plate basis as described for Experiment 1.

For both experiments the Nextflow workflow automatically generated tab-separated files of log2-transformed intensities from each Skyline file created (see Figure 1c). These files had data organized with rows corresponding to features (protein groups or modified peptide precursors) and columns corresponding to individual samples and were used as input for further data analysis as depicted in (see Figure 1d).

### Imputing Retention Time Boundaries for Missing Peaks using Skyline

During import of the DIA-NN search results into Skyline, we enabled Skyline’s peak boundary imputation (”Impute boundaries for missing peaks” function) to recover integration boundaries for peptides that DIA-NN identified in some replicates but not others. For each peptide, Skyline took the median of its observed retention times across the DIA-NN results, producing a per-peptide reference axis independent of any individual acquisition (Figure 2-1). For each replicate, Skyline then fit a LOWESS regression mapping that replicate’s observed retention times onto this shared reference, using every peptide identified in the replicate as an anchor point (Figure 2-2). For each peptide in each replicate where DIA-NN did not supply peak boundaries, Skyline selected a donor replicate, the one in which that peptide had received the best DIA-NN identification score, and transferred its peak boundaries into the replicate with missing bounds (Figure 2-3). The transfer was performed by projecting the donor’s start and end times through the donor replicate’s LOWESS fit into the shared reference axis and then back out through the recipient replicate’s LOWESS fit into the recipient’s own observed-retention-time axis. The resulting start and end times were applied directly to the recipient chromatogram as the integration window, so that each imputed peptide was integrated over a retention-time window consistent with its elution position in the replicate in which it had been most confidently identified. When multiple replicates tied on the best score for a peptide, their transferred boundaries were combined by taking the median rather than selecting a single donor. Because this procedure was carried out automatically during chromatogram import, the resulting Skyline document contained an integration value for every peptide in every replicate, including those for which the upstream DIA-NN search had returned no retention-time information.

**Figure 2.**
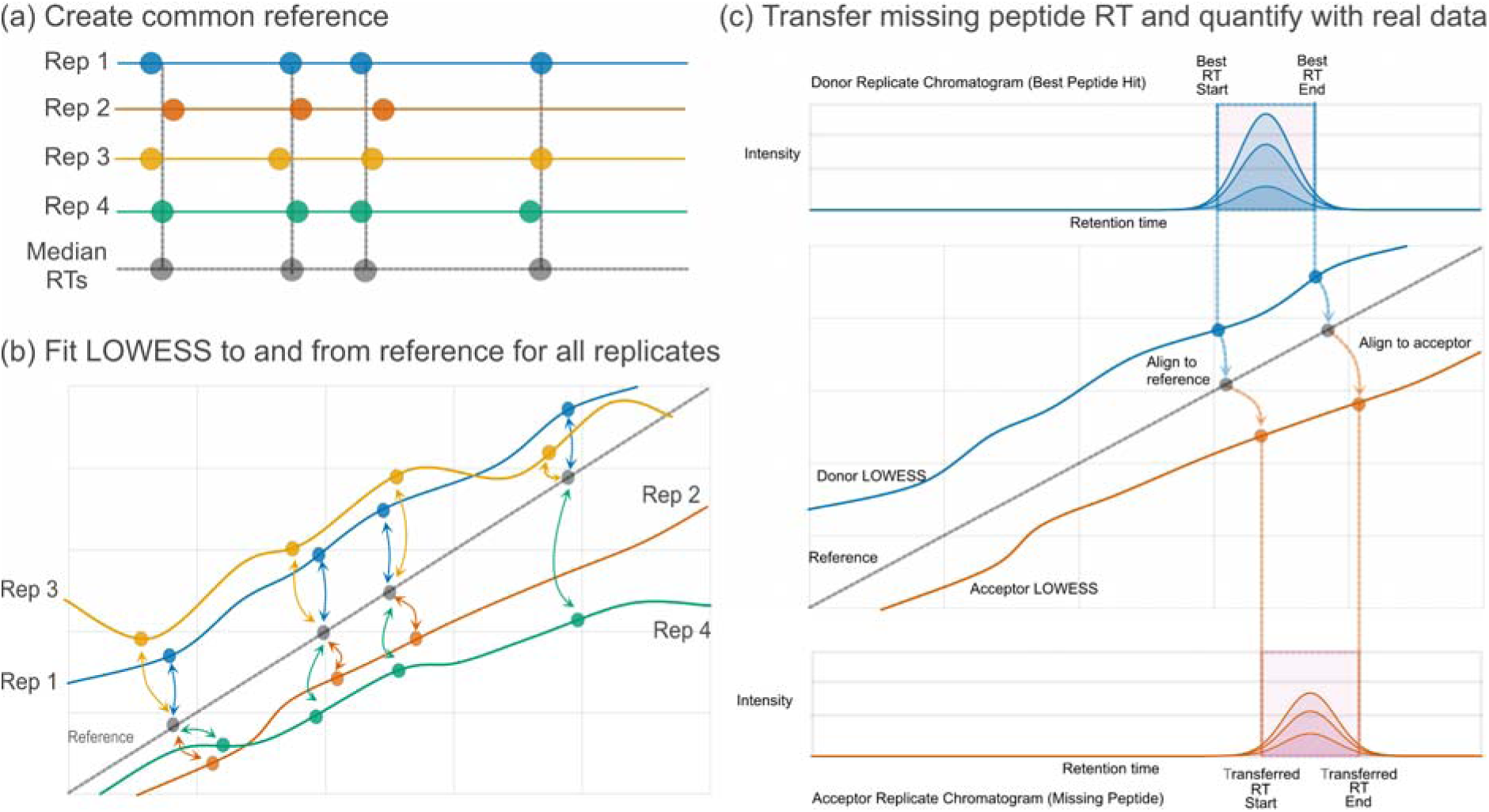
Skyline retention time imputation and signal recovery algorithm. (a) First, a shared reference list of retention times for each identified peptide across all replicates is constructed using the median of retention times observed for each peptide across all replicates. (b) For every replicate, a LOWESS regression is fit to and from the reference based on the observed retention times for peptides in each replicate. (c) For each missing peptide in each replicate, retention times are imputed by first translating the observed retention time for that peptide from the replicate with the best score for that peptide to the reference (using the LOWESS fit from panel b) then translating the reference retention time to the replicate missing the peptide. The abundance of the missing peptide can then be calculated by integrating the signal between the transferred retention time boundaries.

**Figure 3.**
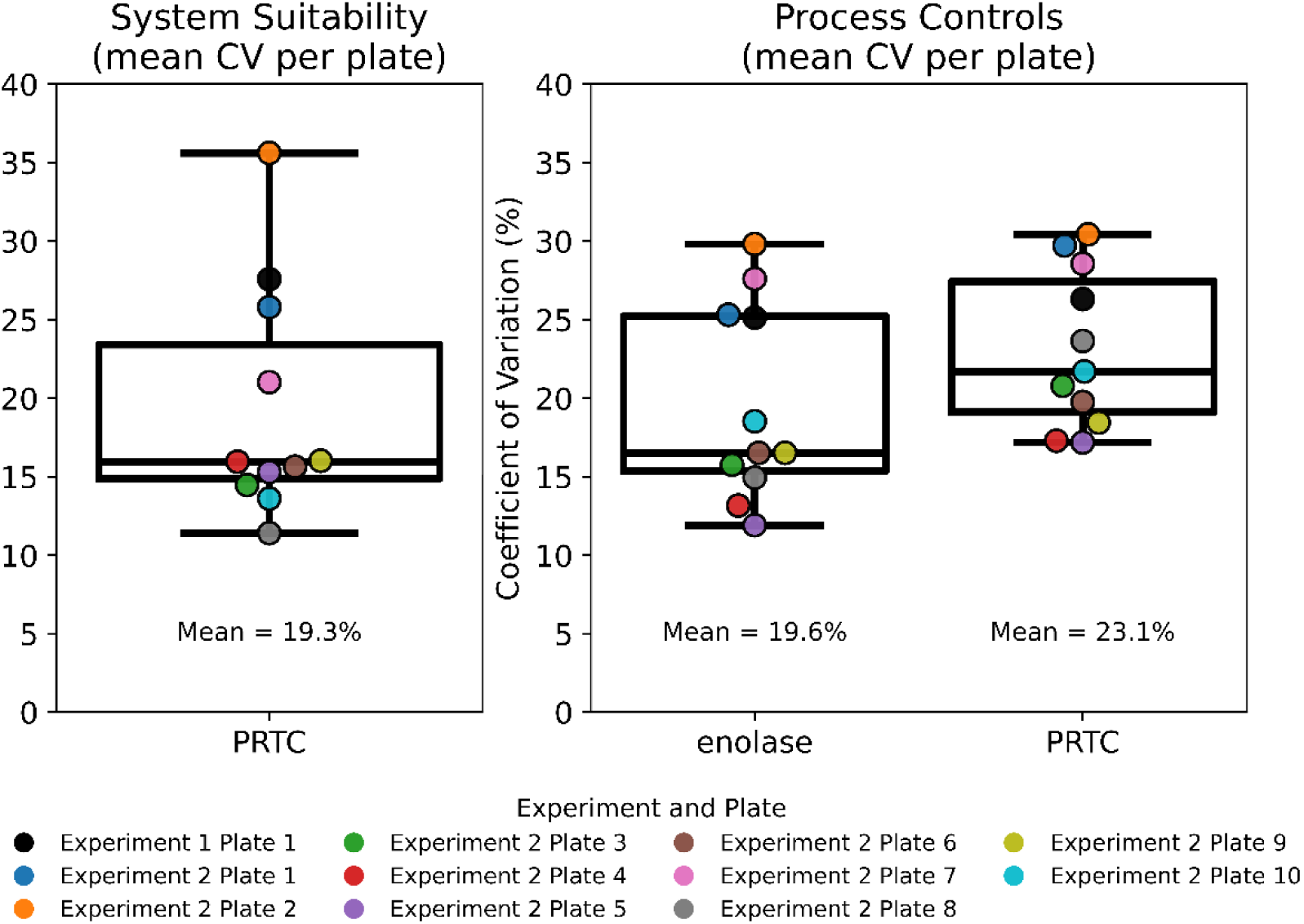
Summary of peptide variance across all plates for process control peptides (yeast enolase), internal QC peptides (process control PRTCs) and external system suitability peptides (system suitability PRTCs). Peptide-level CVs were first calculated for each peptide using peak area values for all replicates within each plate. For each plate, mean CV values were then calculated across all peptides matching either enolase or PRTC peptides. The left panel shows sample independent system suitability injections, which consisted of a tryptic digest of BSA supplemented with PRTC peptides. The right panel shows process control samples, including both enolase an PRTC peptides, which were added to the experimental samples at different stages during sample workup. Each point represents the mean CV for a single plate, with colors indicating the experiment and plate of origin. Boxplots summarize the distribution of plate-level mean CV values across all plates within each category. Text labels within each panel indicate the overall mean CV across all plates for that category.

### Data Analysis and Statistical Methods

#### Normalization and Batch Correction

Protein- and precursor-level abundance matrices were produced as described above. Sample-wise normalization was performed using the pronoms Python package (https://github.com/mriffle/pronoms) for median normalization using the ‘MedianNormalizer’ function. Briefly, log2 intensities were exponentiated to the linear scale and divided by each sample’s median. The mean of the original sample medians across all samples was calculated and the median-scaled values were multiplied by that global mean-of-medians so the normalized data remained on roughly the original intensity scale. Normalized data were then re-log2-transformed. For Experiment 2, following normalization, plate-to-plate technical variation was removed by empirical Bayes batch correction using ComBat^26^ as implemented in the ‘pycombat’ Python package,^27^ with the plate identifier as the batch variable and the full 936-sample matrix passed through correction in a single call before any downstream sample filtering. All array operations were performed using NumPy^28^ and pandas.^29^ For Experiment 1, batch correction was not performed as the entire experiment consisted of a single batch.

#### Principal Component Analysis (PCA)

Principal component analyses (PCA) were performed using un-normalized, median normalized, or median normalized batch corrected precursor or protein level log2 abundance data as produced by our workflow (see Normalization and Batch Correction). Features were standardized to zero mean and unit variance using scikit-learn’s ‘StandardScaler’^30^ and decomposed via ‘PCA’ retaining the first four components. Figures show scatter plots of PC1 vs. PC2 with axis labels reporting the percent variance explained by each component, and each scatter is flanked by marginal panels above and to the right that visualize how a chosen sample annotation distributes along the corresponding principal axis. For categorical annotations, samples are colored by group and the marginal panels show per-group kernel density estimates of the principal component scores using SciPy’s ‘gaussian_kde’;^31^ the per-axis test for a difference in distribution between groups is a two-sided Mann-Whitney U test (‘scipy.stats.mannwhitneyu’) when exactly two groups are present, and a Kruskal-Wallis H test (‘scipy.stats.kruskal’) when there are three or more, with the resulting p-value annotated on each marginal panel. For continuous annotations, samples are colored on a sequential colormap and the marginal panels show a scatter of the principal component score against the annotation value overlaid with a simple linear regression line fitted via ‘scipy.stats.linregress’; the surrounding shaded band is the 95% confidence interval for the *mean response* of the regression, computed analytically from the residual mean square as ‘SE_mean(x*) = sqrt(MSE * (1/n + (x* - mean_x)^2 / Sxx))’ and scaled by the two-sided t critical value with ‘n - 2’ degrees of freedom, matching the convention used by seaborn’s ‘regplot’. The slope and the corresponding two-sided p-value for the null hypothesis that the slope equals zero are extracted from ‘linregress’ and annotated on each marginal panel. All PCA plotting was implemented using Matplotlib.^32^

#### Boruta Feature Selection

To identify proteins whose abundance was informative for dose rate in Experiment 2, we applied the Boruta all-relevant feature selection algorithm as implemented in the ‘BorutaPy’ Python package (https://github.com/scikit-learn-contrib/boruta_py).^33^ The input to Boruta was the normalized, batch-corrected log2 protein abundance matrix (see Normalization and Batch Correction) restricted to the 700 experimental samples (‘Sample_Type == “Skin Punch”’) and split by Focus Area (FA1 or FA2). Boruta used a ‘RandomForestClassifier’ with balanced class weights from scikit-learn.^30^ At each iteration Boruta constructs “shadow” features by randomly permuting each original feature, fits a random forest on the combined original-plus-shadow matrix, and tests via a two-sided binomial test whether each original feature’s importance significantly exceeds the maximum importance observed across the shadow features; features are iteratively classified as Confirmed, Tentative, or Rejected. Any missing values were set to zero before fitting.

#### Classification

Binary classification of samples was performed using elastic-net regularized logistic regression on the normalized, batch-corrected log2 abundance matrix (see Normalization and Batch Correction). The input feature matrix was filtered to the relevant sample subset for each comparison, and any residual missing values were set to zero before fitting. The classifier was implemented as a scikit-learn ‘Pipeline’^30^ chaining a ‘StandardScaler’ to a ‘LogisticRegression’ estimator with the SAGA solver, an elastic-net penalty, and ‘class_weight=”balanced”’ to address class imbalance. Hyperparameters, the inverse regularization strength ‘C’ and the L1/L2 mixing ratio ‘l1_ratio’, were selected via stratified k-fold cross-validated grid search using mean area under the receiver-operating-characteristic curve (AUC) as the scoring metric. To reduce the chance of selecting an unstable single-cell optimum, grid scores were smoothed using a von Neumann neighborhood mean (each cell averaged with its immediate ±1 neighbors along both grid axes), and the smoothed maximum mean AUC was chosen as the optimal configuration; tuning results were cached to disk for reproducibility. The chosen hyperparameters were then evaluated on the same data using repeated stratified k-fold cross-validation (5 folds × 5 repeats = 25 folds), with the ‘StandardScaler’ re-fit on each training fold to prevent leakage of test-fold statistics into the held-out predictions. The output of each run is a mean ROC curve with a ±1 standard deviation band, the cross-validated AUC, and balanced accuracy (which accounts for class imbalance), together with a heatmap of the cross-validated AUC across the ‘C’ × ‘l1_ratio’ grid. All cross-validation, scoring, and ROC computation routines were taken from scikit-learn, and CPU-bound grid cells were evaluated in parallel using joblib.

#### Regression

Continuous outcomes were modeled from proteomic abundance data using elastic-net regularized linear regression on the normalized, batch-corrected log2 abundance matrix (see Normalization and Batch Correction). The input feature matrix was filtered to the relevant sample subset for each analysis, and any residual missing values were set to zero before fitting. The regressor was implemented as a scikit-learn^30^ ‘Pipeline’ chaining a ‘StandardScaler’ to an ‘ElasticNet’ estimator with ‘selection=”random”’ and an intercept term. Hyperparameters, the regularization strength ‘alpha’ and the L1/L2 mixing ratio ‘l1_ratio’, were selected via repeated k-fold cross-validated grid search using mean absolute error (MAE) as the scoring metric. To reduce the chance of selecting an unstable single-cell optimum, grid scores were smoothed using a von Neumann neighborhood mean (each cell averaged with its immediate ±1 neighbors along both grid axes), and the smoothed minimum mean MAE was chosen as the optimal configuration; tuning results were cached to disk for reproducibility. The chosen hyperparameters were then evaluated on the same data using repeated k-fold cross-validation (10 folds × 5 repeats = 50 folds), with the ‘StandardScaler’ re-fit on each training fold to prevent leakage of test-fold statistics into the held-out predictions; predictions were optionally clipped to a non-negative floor (and, where appropriate, an upper ceiling) to enforce physical plausibility before scoring. The output of each run is a per-sample averaged true-vs-predicted scatter plot with a fitted regression line, MAE ± standard deviation, and prediction R² ± standard deviation, together with a heatmap of the cross-validated MAE across the ‘alpha’ × ‘l1_ratio’ grid. All cross-validation, scoring, and grid search routines were taken from scikit-learn, and array operations used NumPy.^28^

#### Generalized Additive Model (GAM)

To identify proteins whose abundance was associated with experimental covariates, we fit a generalized additive model^34^ to each feature independently using the pyGAM Python package.^35^ The input was the normalized, batch-corrected log2 abundance matrix (see Normalization and Batch Correction) restricted to the relevant sample subset, with any residual missing values set to zero before fitting. Each per-feature model expressed log2 abundance as a sum of penalized cubic spline smooth terms (‘s()’, with 20 basis functions and the default penalty) for continuous covariates whose response was expected to be nonlinear, and unpenalized linear terms (‘l()’) for binary or categorical covariates whose effect was to be tested directly via a Wald-type coefficient. Interactions between a continuous and a binary covariate were modeled with a by-variable smooth (‘s(x, by=z)’), which captures the additional response contributed by one group beyond a shared baseline trajectory; an interpretable effect size for such terms was computed as the average slope of the by-variable smooth across the observed range of the continuous covariate using pyGAM’s ‘partial_dependence’ method. Coefficient estimates and p-values for each model term were extracted from the fitted GAM, and multiple-testing correction was performed independently per term across all features using the Benjamini-Hochberg false discovery rate procedure^36^ as implemented in the ‘multipletests’ function of statsmodels.^37^ Per-feature fits were parallelized across CPU cores using joblib, and any individual fits that failed to converge were recorded as missing rather than dropped.

#### Data Records

The complete datasets for Experiment 1 and Experiment 2 are available via Panorama Public^38^ (https://panoramaweb.org/teirex-2a-ldxr-mouse-pelt.url) and were assigned the ProteomeXchange^39^ ID: PXD078423 (https://doi.org/10.6069/5ey2-j220). Experimental data are provided at multiple levels of processing, from raw MS data (Level 0) to normalized peptide-and protein-level quantities suitable for downstream analysis (Levels 3a and 3b, respectively), as summarized in Figure 1e. **Level 0** consists of raw data. For Experiment 1 vendor raw files and mzML files are provided. The mzML files were generated using ProteoWizard (3.0.25168) with demultiplexing applied during conversion to allow direct processing by most proteomics analysis software. For Experiment 2 only vendor raw files are provided as data were acquired without overlapping windows and therefore do not require demultiplexing prior to analysis. **Level 1** comprises zipped Skyline documents (one per 96-well plate). **Level 2** contains unnormalized peptide- and protein-level data, and **Level 3** contains normalized (Experiment 1) or normalized plus normalized and batch-corrected (Experiment 2) peptide- and protein-level data.

Skyline documents and raw data for system suitability and process controls are available in the corresponding sections of the Panorama Public site. Process control data are contained within the sample raw files, as controls were spiked into each sample. System suitability data were acquired separately using dedicated runs designed to assess LC-MS/MS performance and are thus provided as separate data files. These sections of the Panorama Public site follow the same framework as for the experimental data except that normalized data is not provided as it is not desirable for system suitability or process control analytes, which are spiked into samples in defined quantities.

### Technical Validation

#### Experimental Design and Internal Controls

The mice used in Experiment 1 of this study were exposed at the University of Washington (UW), while the mouse pelts used for the experimental samples in Experiment 2 were produced at Lawrence Berkeley National Laboratory (LBNL). The X-ray sources at each facility were calibrated according to strict standards (see X-ray exposure section above). In addition, inter-facility dose harmonization measurements were made using EBT3 radiochromic film. Results showed cross-site dose agreement within 12% of expected values, with an average deviation of approximately 6% below the nominal dose (data not shown).

For both experiments, animals were housed in groups and randomly assigned to experimental groups by cage. Mouse pelt samples were collected immediately following euthanasia. Mice were shaved, pelts were dissected and skin punches were collected and transferred into sterile 1.5 mL microcentrifuge tubes, frozen on dry ice and stored at −80°C in one continuous protocol.

Prior to proteomic analysis, all study samples were randomized. This randomized order was preserved throughout sample preparation, digestion, and MS data acquisition and is recorded in the metadata of both experiments: see Supplementary Tables S1 and S2 (Supplementary_Table_S1.xlsx and Supplementary_Table_S2.xlsx) for metadata headings and https://panoramaweb.org/teirex-2a-ldxr-mouse-pelt.url for complete metadata for both experiments. Experiment 1 was a single batch and the “WellPosition” column in the metadata represents this randomized order. For Experiment 2 the metadata has “Plate” and “Well_Position” columns. Samples were prepared and run as individual plates starting with plate P01 and proceeding through P10 in that order. Within each plate the samples were ordered by Well_Position.

A system of process controls and system suitability standards, previously described by Tsantilas et al, 2024,^14^ was used to ensure high quality of the entire dataset. These were added at different stages of sample preparation as described in the Methods section and illustrated in Figure 1b. All experimental samples included Pierce Retention Time Calibration (PRTC) mixture as an internal process control. These peptides enable monitoring of chromatographic retention time and MS signal intensity stability across runs independent of protein digestion efficiency. In addition, intact yeast enolase protein was added to all samples prior to digestion to capture variability introduced during sample preparation, digestion, chromatography, and MS data acquisition. Finally, sample-independent system suitability injections consisting of a tryptic digest of BSA supplemented with PRTCs were also interspersed throughout the experimental sample acquisition sequence, every 6 to 12 sample injections. These sample-independent controls enable longitudinal monitoring of LC–MS/MS performance independent of sample composition.

The mean CV of the 15 PRTC peptide peak areas in the sample independent system suitability injections was 19% across all 11 plates acquired across both experiments (Figure 3, left panel). A similar mean CV of 23% was observed (Figure 3, right panel) for PRTCs measured within the experimental study samples, a total of 1,032 experimental sample injections, indicating that the autosampler, chromatography and MS instrument performance was within specifications during acquisition of sample data. The mean CV of the 4 enolase peptides monitored across all runs was 20% indicating that sample workup and digestion did not add significantly to run-to-run variation.

#### Peptide and Protein Detections in Mouse Pelt Samples

Using a 1% false discovery rate (FDR) cut-off, between 68,657 and 114,740 unique precursors were detected by DIA-NN across all 96 samples in Experiment 1. These precursors were mapped to between 6,129 and 8,323 protein groups by DIA-NN. Within Experiment 2 between 89,779 and 151,581 unique precursors were detected by DIA-NN across all 936 samples. These precursors were mapped to between 7,380 and 9,574 protein groups by DIA-NN (Figure 4). Experiment 1 was performed on a different mass spectrometer (a Thermo Orbitrap Eclipse) to Experiment 2 (a Thermo Astral), which likely accounts for the different number of precursor and protein detections.

**Figure 4.**
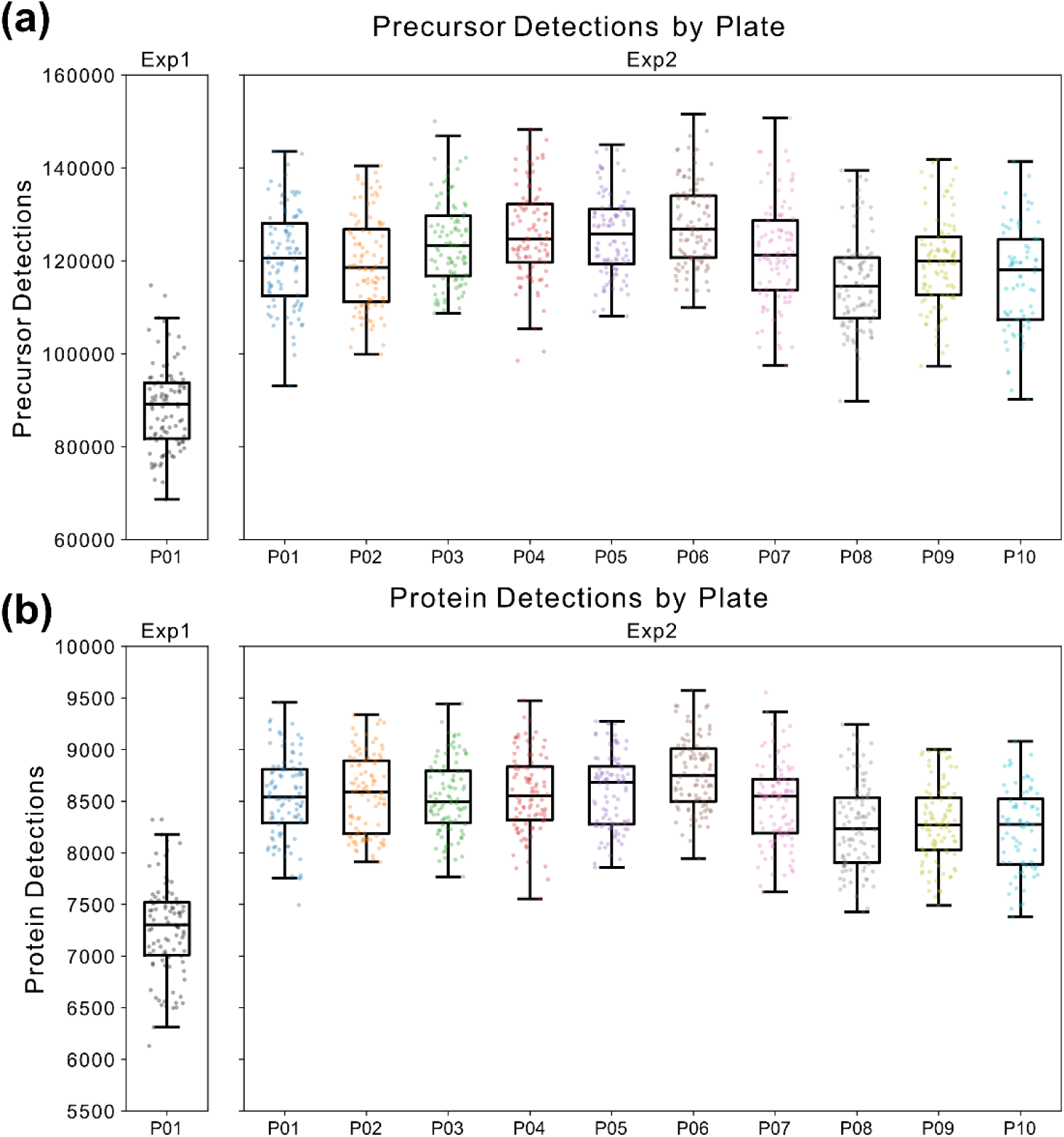
Box and Whisker plots of precursor and protein detections across all plates of Experiment 1 and Experiment 2. (a) The number of precursors identified by DIA-NN in each LC-MS/MS injection. (b) The number of proteins identified by DIA-NN identified per LC-MS/MS injection. Samples are grouped by Experiment Number (Exp1 or Exp2) and plate number (P01 through P10). Data is presented at 1% FDR.

In the current study, precursor peptide identifications and retention time boundaries were initially generated independently for each sample using DIA-NN. These results were then imported into Skyline through our Nextflow workflow (Figure 1c), where identifications from all samples within an experiment were combined to generate a common set of precursor peptide targets and retention time boundaries across the entire dataset. Skyline’s “Impute boundaries for missing peaks” function was then used to align retention times across samples and define peak boundaries for targets that were not initially detected in a given sample. As a result, in our fully processed datasets, the same set of precursor peptide targets was quantified in every sample, allowing either precursor signal or background signal to be extracted from the expected retention time window even when no peptide peak was originally identified. The details of this are described in the Methods section under the subheading Imputing Retention Time Boundaries for Missing Peaks using Skyline.

Using this approach, Experiment 1 yielded quantification of 123,771 unique precursors, 100,311 unique peptides, and 9,562 unique proteins. Experiment 2 yielded quantification of 154,911 unique precursors, 131,109 unique peptides, and 10,568 unique proteins.

#### Reproducibility and Verification

Previous experiments using similar mouse pelt production and lysis methods indicated that protein concentrations in the resulting lysates were relatively consistent and fell within a range that could be effectively accommodated by downstream computational normalization strategies.^40^ Therefore, rather than measuring protein concentration for each sample and adjusting the amount of lysate added to each digest, a fixed lysate volume was used for all samples. For the current study, this volume was set at 50 µL per sample, which, based on previous measurements from similar lysates, was estimated to yield prepared digests containing approximately 1 µg of total protein per 3 µL.

As described in the Normalization and Batch Correction section, median normalization was applied to protein- and precursor-level abundance matrices. This was done to compensate for unavoidable variation introduced during sample processing and MS data acquisition, as well as differences in protein input resulting from the use of a fixed lysate volume for all digests regardless of individual sample protein concentration. For Experiment 2, following normalization, plate-to-plate technical variation was removed by empirical Bayes batch correction using ComBat.

The distribution of protein peak areas in each sample is shown in Figure 5, left panels, before and after median normalization for Experiments 1 and 2 and additionally after batch correction for Experiment 2, which constituted 10 plates, each of which was considered a batch. Our experimental design incorporated sample randomization prior to proteomic sample preparation and this randomized order was kept throughout MS data acquisition (see Methods section). Samples representing different radiation doses, dose rates, harvest times, and other experimental variables were distributed across plates and acquisition batches rather than being grouped together. This reduced the potential for confounding between biological effects and technical variation and enabled more effective downstream normalization and batch correction.

**Figure 5.**
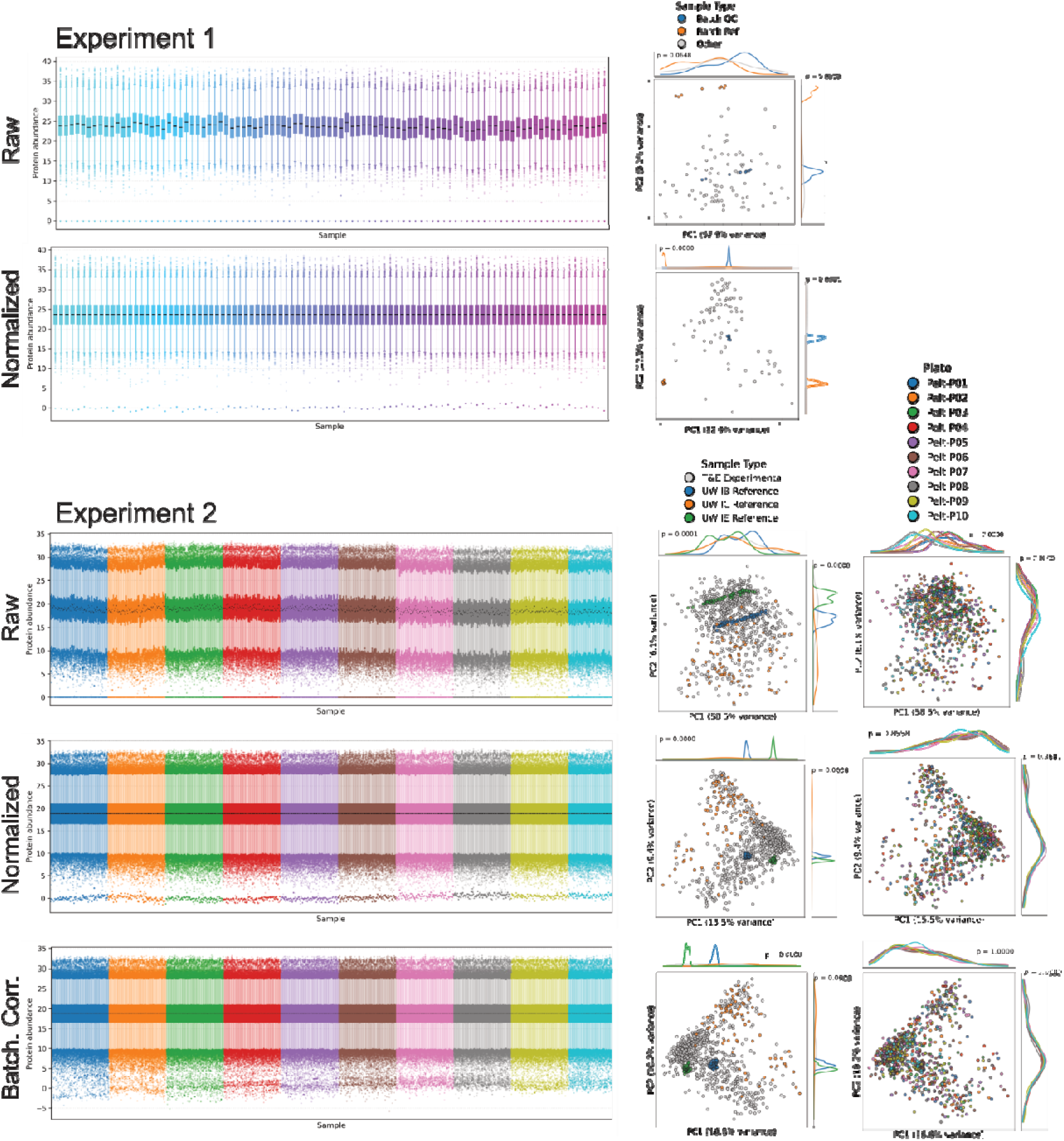
Visualizing per-search protein abundance distribution and the effects of normalization and batch correction. The first column shows the distribution of log2 protein abundances in all searches for Experiments and 2, sorted by acquisition order. For Experiment 1, a raw (unnormalized) row and median-normalized row are shown. For Experiment 2, a raw, median-normalized, and batch-corrected row are shown. Experiment 1 is colore according to acquisition order, and Experiment 2 is colored by 96-well plate and is sorted by acquisition order. The second column shows the results of a principal component analysis (PCA) based on sample for the raw, normalized, and batch-corrected (Experiment 2) protein abundances. For Experiment 1, blue points are batch QC, orange are batch reference, and green are experimental data. For Experiment 2, orange points are the inter-batch reference, red are the inter-experiment reference, green are the internal control (embedded Experiment 1), and blue are the experimental data. The third column (Experiment 2 only) is the same PCA analysis but colored by 96-well plate. The marginal density plots along the top and right side of the PCA plots indicate the density along that principal component for the category of interest, colored using the same colors as the PCA points. P-values shown on the marginal panels indicate the strength of evidence against the null hypothesis of no distributional difference (categorical) or no association (continuous) along each principal component; see Methods for details.

Principal component analysis (PCA) was used to visualize high-level structure in the proteomic abundance data and to assess how that structure related to sample annotations. Figure 5, middle panels, show PCAs colored by sample type. Experiments 1 and 2 each incorporated 2 independent reference samples per row of each 96-well plate. The replicates of these reference samples are separated in the PCAs of unnormalized data but form tight clusters after median normalization indicating that the normalization is compensating for experimental variation.

Experiment 2 additionally incorporated a set of internal control reference samples consisting of all 80 samples from Experiment 1. These samples were re-digested and reanalyzed within Experiment 2 and randomly interspersed across all 10 plates. They were included as positive controls to enable evaluation of the performance of the larger Experiment 2 dataset in capturing radiation-associated proteomic phenotypes. Experiment 2 PCAs colored by sample type show these reference samples are distributed within the spread of the 700 experimental samples regardless of normalization or batch correction. This indicates the internal control samples are similar to the 700 experimental samples, which were produced in a different facility, and suggests that the Experiment 1 samples are a useful internal control for Experiment 2. The fact the samples retain a diffuse spread after normalization and batch correction is expected as each of these reference samples are from different animals with different exposure levels. This spread represents the biological variability of interest.

Figure 5, Experiment 2 right panels, show PCA plots colored by plate number. Prior to normalization, samples separate strongly by plate (p = 0.0), consistent with technical variation introduced because each plate was prepared and acquired separately. Median normalization reduces this plate-associated structure, while batch correction removes it almost entirely, resulting in no significant association between plate number and the major principal component axes (PC1 and PC2; p = 1.0).

An analysis of CVs of the independent reference samples incorporated into each experiment showed median CVs of 30 and 25% for Experiment 1, and 30 and 31% for Experiment 2 before normalization. These CVs, impacted by technical variation between multiple digests of the same reference pool across different rows and plates within an experiment, were reduced to 14 and 11% by median normalization in Experiment 1, and to 14 and 15% in Experiment 2 by normalization and batch correction (Figure 6 panels a, b, d and e).

**Figure 6.**
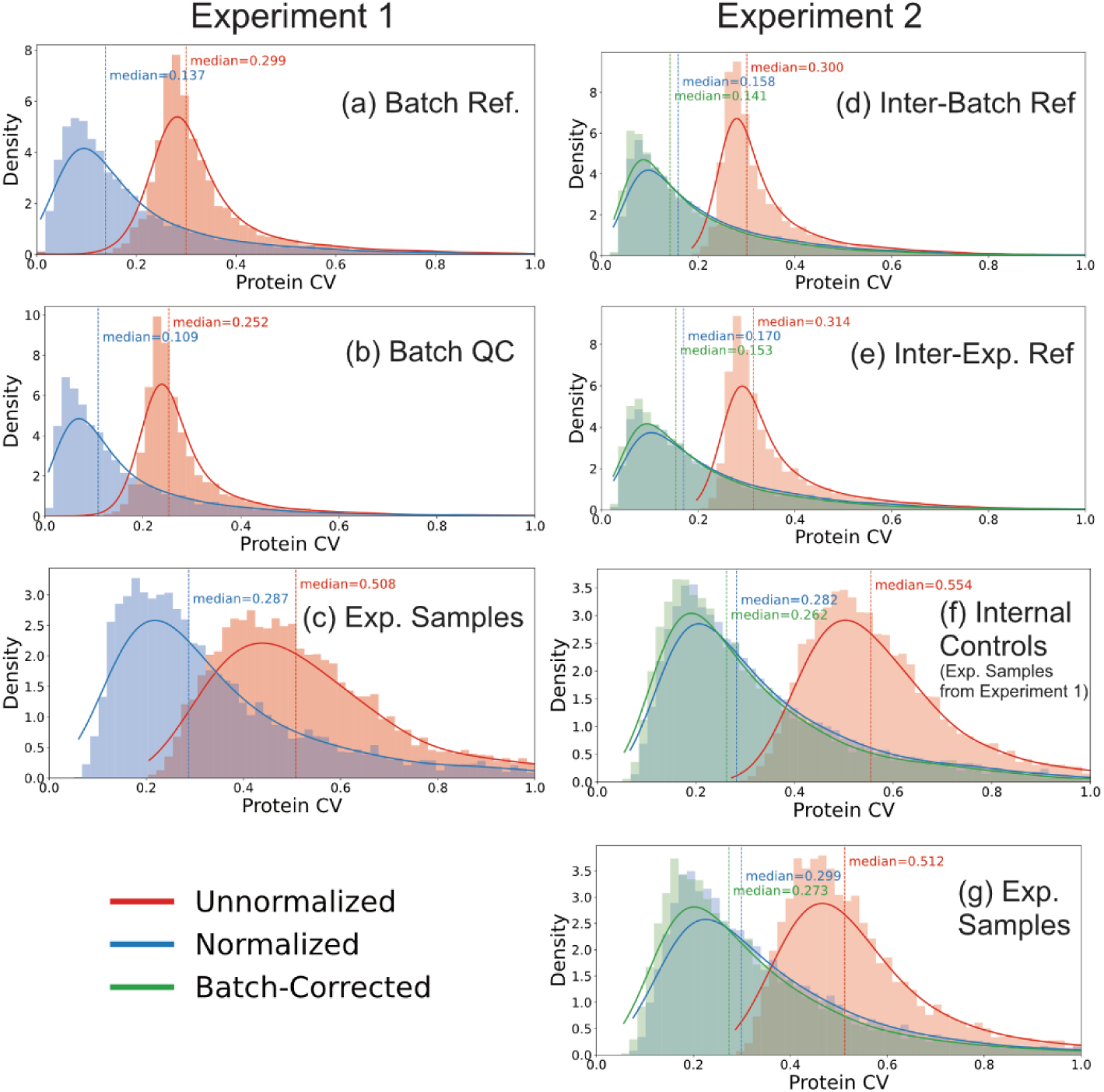
The effects of normalization and batch correction on protein-level coefficients of variation (CV). The coefficients of variation for the non-logged abundances for all proteins in Experiments 1 and 2 were calculated and their frequency plotted for raw (red), median-normalized (blue), and median normalized plus batch corrected (green, Experiment 2 only) data. For Experiment 1 (first column) the top plot shows CVs for proteins in the batch reference samples, the middle plot shows CVs in the batch QC samples, and the bottom plot shows CVs in experimental samples. For Experiment 2 (second column), the top plot shows CVs in the inter-batch reference samples, the second plot shows CVs in the inter-experiment reference samples, the third plot shows CVs in the internal controls (embedded samples from Experiment 1). The bottom plot shows CVs in the experimental data. The median of the distributions of the CVs for raw, normalized, and batch-corrected data are shown as text with the same color as their respective distribution.

The 80 experimental samples in Experiment 1 are represented in Figure 6 panel c and show a median CV of 51% before normalization and 29% after normalization. CVs in these samples are expected to be higher than for the reference samples as the experimental samples are from different individual animals exposed to different treatments. These samples should therefore contain biological variability of interest. These same 80 samples were included as internal controls in Experiment 2, randomly distributed across all 10 plates. In Experiment 2 these samples were named “internal controls” (IC), and are depicted in Figure 6 panel f. They show a median CV of 55% before normalization, 28% after normalization and 26% after normalization and batch correction. The fact that the CVs of these same samples were similar when embedded within Experiment 2, versus run alone in Experiment 1 illustrates our ability to run 10 full plates constituting 936 total samples, while maintaining good reproducibility and computationally accounting for any technical variability and potential batch effects that might be expected from such a large experiment.

The 700 experimental samples analyzed in Experiment 2 resulted in median CVs of 51% before normalization, 30% after normalization and 27% after normalization and batch correction (Figure 6 panel g). These CVs are similar to the internal controls and show the expected variability for individual animals exposed to differing treatments and containing biological variability of interest.

Taken together the data presented in Figures 5 and 6 show the high quality of the current datasets, the value of the reference samples and the internal controls and the fact that biological variability was preserved after normalization and batch correction.

### Usage Notes

We provide peptide- and protein-level data from mouse dorsal skin punch samples collected following controlled X-ray exposures spanning a range of doses, dose rates, and post-exposure time points. This dataset enables investigation of proteomic changes associated with radiation exposure and supports evaluation of how these changes vary with dose, dose rate, and time after exposure. The scale of the dataset, together with its extensive reference samples and technical controls, also makes it well suited for the development and benchmarking of normalization methods, biomarker discovery strategies, and machine learning approaches for radiation biodosimetry.

#### Use Case 1: Biomarker Discovery for Response to Dose Rate

The data may be used to find protein biomarkers that respond specifically to radiation dose *rate*. This could reveal how tissue distinguishes a brief, intense radiation insult from the same total dose delivered more slowly. Dose rate changes the biology of exposure by altering the balance between damage induction and repair during irradiation, and it can also shift cell-state programs such as checkpoint activation and cell-cycle redistribution.^41^ A model built using dose-rate-dependent protein feature lists could help distinguish acute from protracted radiation exposures, potentially informing predictions of injury severity and treatment decisions. Boruta^33^ is a powerful feature selection algorithm built around the Random Forest^42^ classifier. Boruta focuses on finding *all relevant features* by finding features that separate classes in the data more consistently than shuffled shadow features. We applied Boruta to Experiment 2 separately for FA1 and FA2, where FA1 included early post-irradiation time points (6–21 days) and FA2 included later time points (90–150 days), to identify proteins that respond specifically to low dose rate (LDR, 3 cGy/min) versus high dose rate (HDR, 28 cGy/min) exposure (Figure 7).

**Figure 7.**
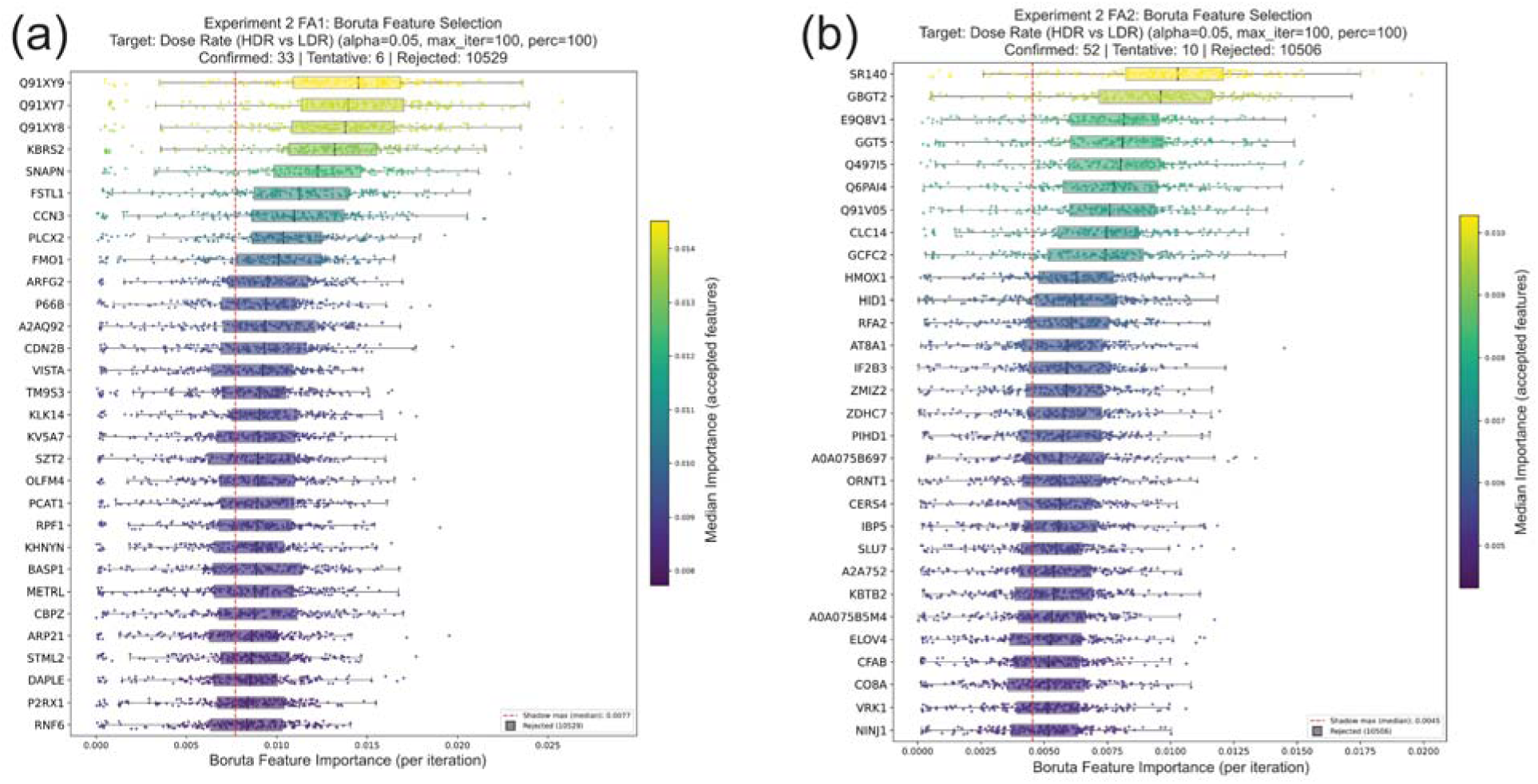
The results of Boruta feature selection for low dose rate (3 cGy/min) versus high dose rate (28 cGy/min) applied to Experiment 2 for (a) FA1 and (b) FA2. The plots show the top thirty proteins ranked by Boruta decision; for each, the horizontal box plot summarizes per-iteration importance values with individual iterations overlaid as jittered points. Box color encodes median importance on a viridis gradient. The vertical dashed red line marks the median per-iteration shadow-maximum importance, which is the effective threshold a protein must exceed to count as a hit. Full data for this plot are provided in Supplementary Table S3 (Supplementary_Table_S3.xlsx).

#### Use Case 2: A Model for Predicting if a Mouse Has Been Exposed to Any Low-Dose Radiation

The data in Experiments 1 and 2 may be used to develop machine learning models to predict if a mouse has been exposed to a low dose (between 5 and 100 cGy) of ionizing radiation using only a skin punch from the mouse within a given number of days from the exposure. This may be useful to develop a proof-of-concept, non-invasive assay to determine if an animal has been near any ionizing radiation source recently. If successful, such a model could, in principle, be developed for other organisms, including humans. The uses for such an assay could include testing for radioactive contamination and exposure near disaster sites or in workplaces where such exposure is possible. For this example model, we developed classifiers with logistic regression with elastic net regularization (see Methods section) to classify controls (0 dose) versus any dose (ranging from 5 to 100 cGy). We developed separate classifiers for Experiment 1 and Experiment 2 (broken out by FA1 and FA2). Each model was built and validated with all experimental data in each set and includes all doses, dose rates, and times since exposure present in each dataset. We evaluated the models using stratified 5-fold cross-validation repeated 5 times (Figure 8). Experiment 1 showed a very strong signal for predicting exposure. Without using a predefined feature list, the classifier achieved an estimated ROC AUC of 0.89 and an estimated accuracy of 0.73. When the analysis was restricted to a predefined set of 92 proteins previously identified as likely to respond to ionizing radiation in mouse pelt tissue (see Supplementary Table S4; Supplementary_Table_S4.xlsx), the estimated ROC AUC decreased to 0.77, although the uncertainty of this estimate increased. In contrast, the estimated classification accuracy increased slightly to 0.74. Classifiers built with Experiment 2 (FA1 and FA2 separately) do contain signal, although it is weaker than that of Experiment 1 (Figure 8). Using the 92-protein feature list improves performance marginally, particularly for FA1, improving the estimated ROC AUC from 0.6 to 0.65 (Figure 8).

**Figure 8:**
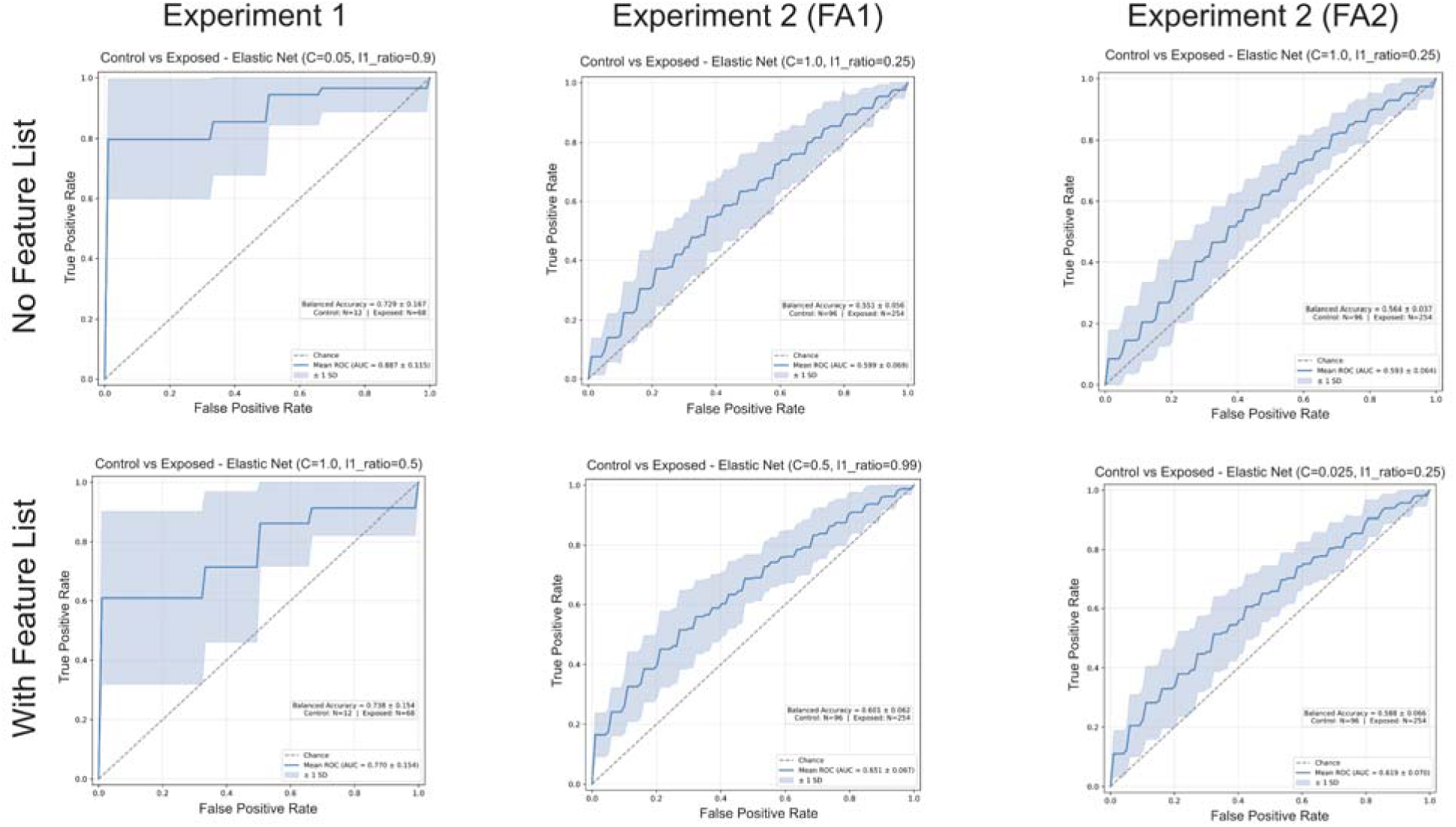
Receiver Operating Characteristic (ROC) curves for exposure classifiers. Each column is an experiment: Experiment 1, Experiment 2 (Focus Area 1), and Experiment 2 (Focus Area 2). The top row is a classifier built using all discovered proteins. The bottom row is a classifier built using only 92 mouse pelt proteins previously found to respond to ionizing radiation. The diagonal dashed line (AUC = 0.5) represents chance-level discrimination, equivalent to random guessing. As AUC approaches 1.0, the model has better ability to rank positive cases above negative cases, with AUC = 1.0 indicating perfect class separation.

If we consider that in Experiment 2, very low doses (under 30 cGy) may be indistinguishable from controls, making the classification task especially difficult. A classifier to distinguish low doses (including controls) from high doses may be more productive. We built classifiers for Experiment 2 (FA1 and FA2) that classified all low doses (0-25 cGy) from higher doses (70-100 cGy). In the case of not using a feature list, both classifiers performed substantially better than classifying 0 versus any dose (Figure 9 versus Figure 8). When using the 92-protein feature list, FA1 saw a substantial gain in terms of both estimated ROC AUC and accuracy, where FA2 saw a detrimental effect (Figure 9). This is likely due to the feature list being generated from a previous experiment with shorter times since exposure coupled with a different proteomic response to ionizing radiation at shorter time points versus longer time points, making the feature list ill-suited to FA2.

**Figure 9:**
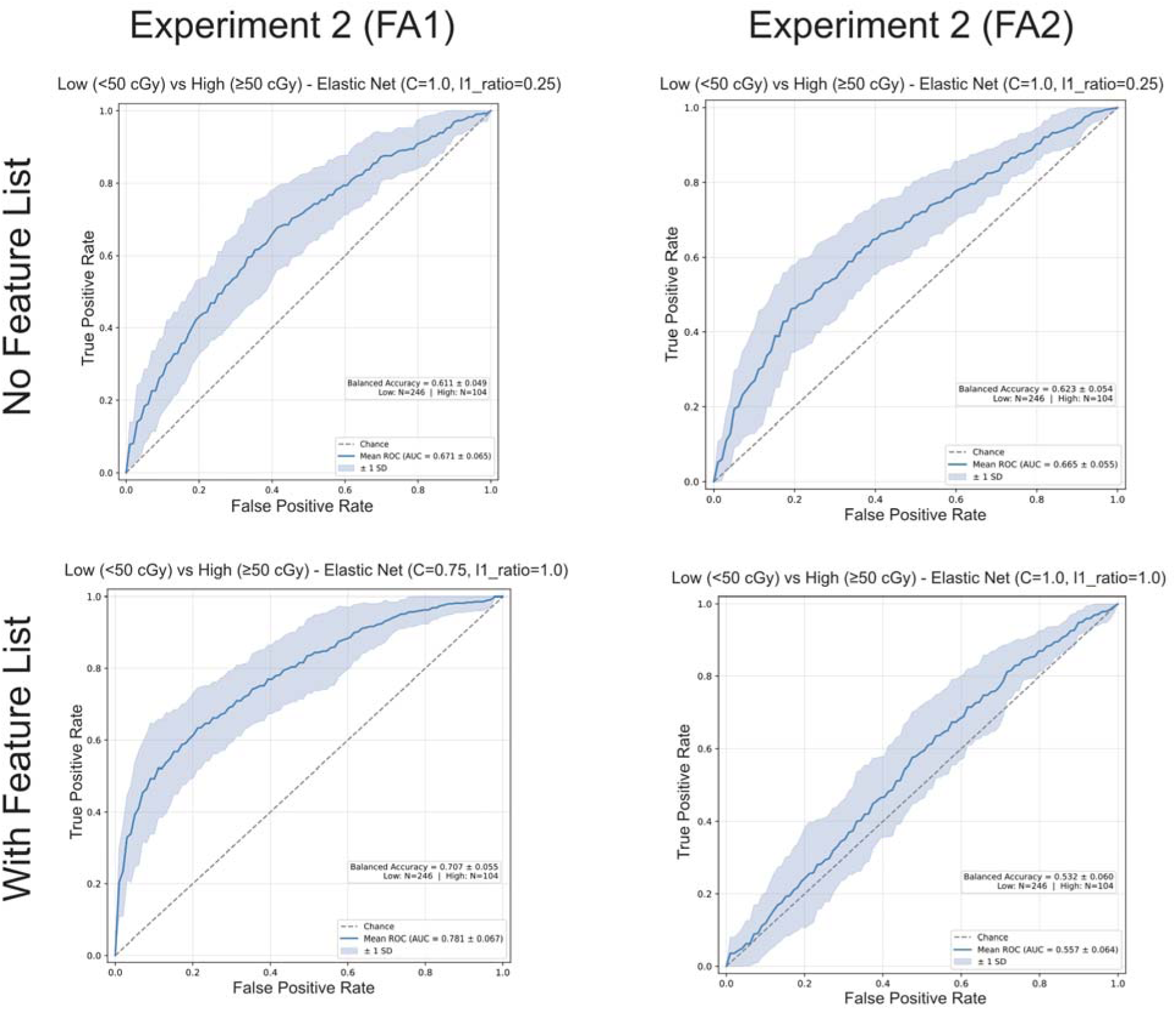
Receiver Operating Characteristic (ROC) curves for low dose (0-25 cGy) versus higher dose (70-100 cGy) classifiers. Each column is an experiment: Experiment 2 (Focus Area 1), and Experiment 2 (Focus Area 2). The top row is a classifier built using all discovered proteins. The bottom row is a classifier built using only 92 mouse pelt proteins previously found to respond to ionizing radiation. The diagonal dashed line (AUC = 0.5) represents chance-level discrimination, equivalent to random guessing. As AUC approaches 1.0, the model has better ability to rank positive cases above negative cases, with AUC = 1.0 indicating perfect class separation.

#### Use Case 3: A Model to Predict the Actual Radiation Dose Delivered to the Mouse

Although the doses are small (ranging from 5 to 100 cGy), it is possible to develop a relatively simple linear regression model to predict dose of X-ray exposure delivered to a mouse using only changes in the proteome abundances in the mouse pelt. Such a model is a valuable proof of concept that it possible to build a relatively non-intrusive assay for assessing the dose of ionizing radiation an individual may have been exposed to days after the exposure so that appropriate action may be taken. In principle, such a model could be developed for other organisms, including humans, and could be a valuable tool for assessing the significance of incidental environmental or workplace exposure to ionizing radiation.

We developed a relatively simple linear regression with elastic net regularization (see Methods section), where we used the abundances of proteins in mouse pelts to predict the dose exposure of the mouse. We built the model using the Experiment 1 samples embedded as a quality control experiment in the larger Experiment 2. The samples include all the doses and dose rates present in Experiment 1. Without using a feature list, the regression model struggled to build an accurate model, likely because regularization struggled to reliably discover the set of proteins with the best response to dose (Figure 10a). However, when we limited the set of proteins to our predefined set of 92 proteins previously identified as likely to respond to ionizing radiation, the model performance improved dramatically, from an estimated MAE of 20.7 cGy to 12.0 cGy (Figure 10b).

**Figure 10:**
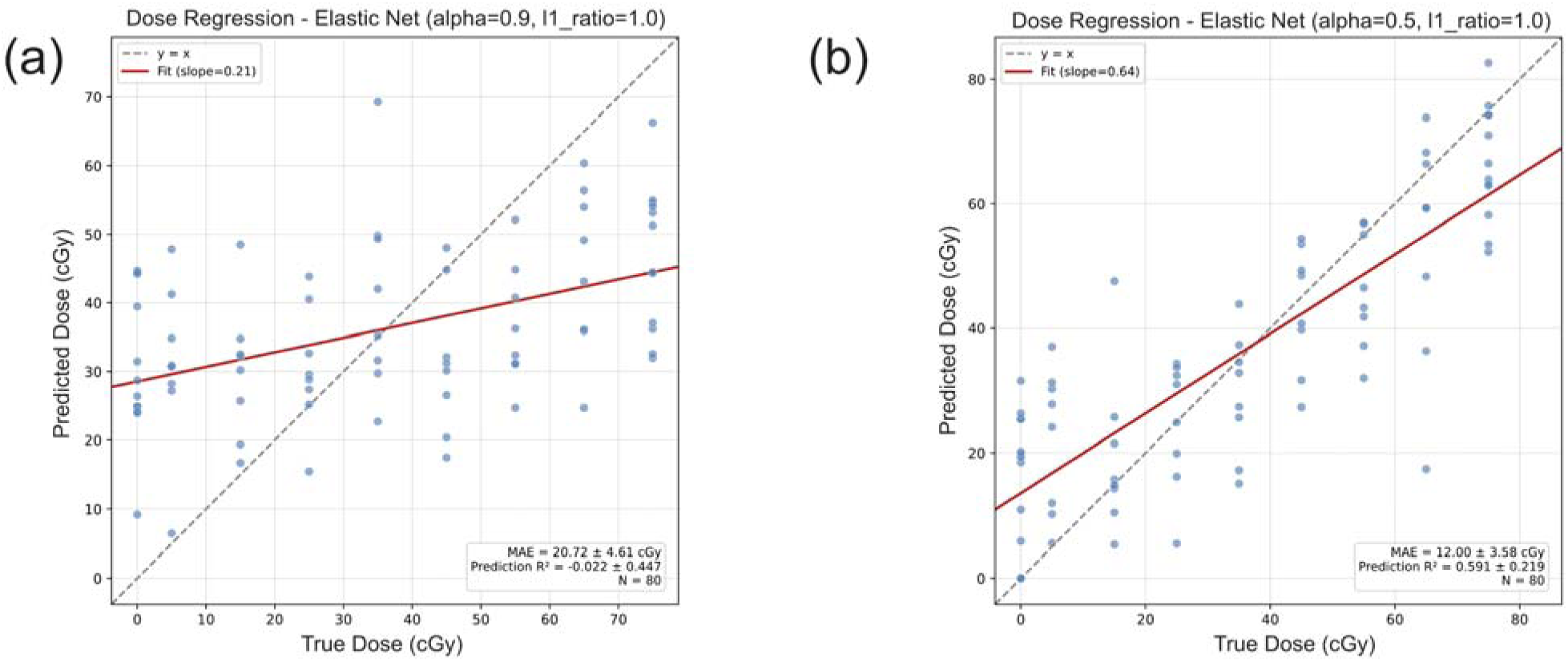
Estimated performance of a linear regression model to predict dose from mouse pelt protein abundances. The displayed results were produced by 10-fold cross-validation repeated 5 times. The scatterplot plots the predicted dose from the test sets from cross-validation versus the true dose. The dashed line indicates perfect predictions, and the red line indicates the fit from the predictions. (a) The result of a model built using all proteins present in the data. (b) The result of a model built using only the 92 proteins found in previous experiments to have a response to ionizing radiation.

#### Use Case 4: Find Protein Targets to Predict Time Since Exposure to Ionizing Radiation

Being able to predict time since low-dose radiation exposure would be valuable because it turns the proteome into a kind of biological clock, helping distinguish recent exposure from later, post-exposure states. That matters for biodosimetry, since the same mouse could show very different molecular patterns depending on when it was sampled after exposure, even at the same dose. The proteins that make such a model work could point to the sequence of biological responses over time, such as early stress signaling, DNA damage response, inflammation, immune activity, tissue remodeling, and recovery. The particular biomarkers used in such a model could provide insight into the kinetics of how skin and associated tissues respond to and recover from irradiation. Finding proteins that may be used as biomarkers to predict time since exposure is confounded by aging of the mouse. Experiment 2 includes controls to help mitigate this, where the mice were euthanized on the same collection day after sham irradiation as real irradiation. The proteins found to respond to time in the sham irradiated mice can be used to help refine the list of candidates to those that depend specifically on being irradiated. To investigate this, we built a Generalized Additive Model (GAM) to build a non-linear model that predicts protein abundance from time since exposure, whether it has been irradiated, and an interaction term for time since exposure and irradiation status (see Methods section). The results of this analysis found 432 proteins with a statistically significant response to time since exposure that was dependent on irradiation status (Figure 11, Supplementary Table S5; Supplementary_Table_S5.xlsx).

**Figure 11.**
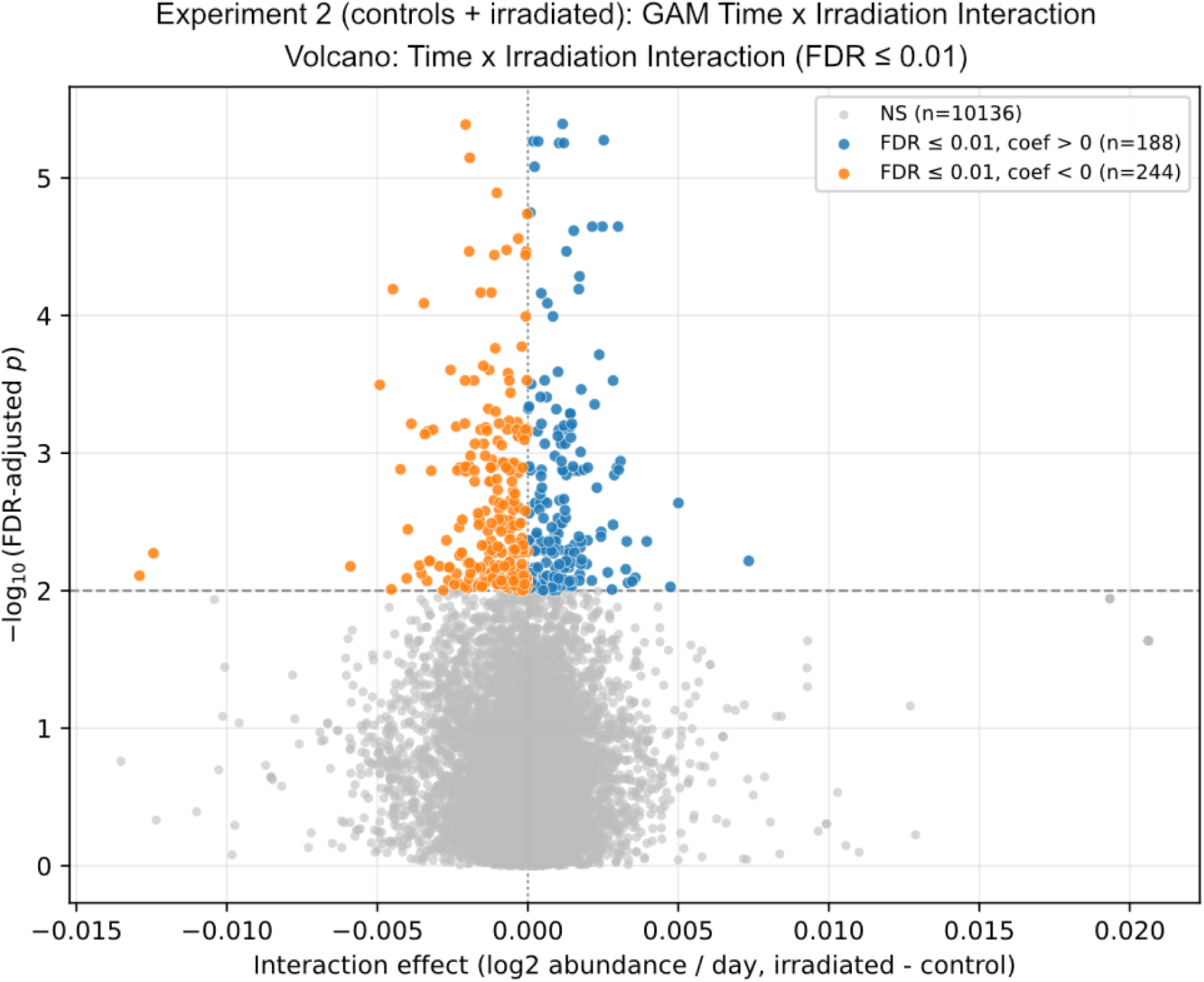
Results of GAM feature finding for proteins with a radiation-dependent response to time since exposure in Experiment 2. The X axis is the average slope of the interaction smooth, which shows how differently a protein’s abundance changes over time in irradiated mice compared to control mice. A positive value (blue dots) means the protein abundances rise faster (or decrease slower) in irradiated mice, and a negative value (orange dots) means the protein abundance drops faster (or rises more slowly) in irradiated mice. The y-axis indicates the −log10 of the B-H adjusted p-value. The gray dots are proteins without a statistically significant adjusted p-value. Full data for this plot are provided in Supplementary Table S5 (Supplementary_Table_S5.xlsx).

#### Use Case 5. Reuse and Reanalysis of the Mass Spectrometry Data for the Development and Benchmarking of Normalization Methods

All peptide- and protein-level data, extracted ion chromatograms, and peptide identifications are available through the Panorama server for reuse and reanalysis. The scale of the current dataset, comprising 1,032 total samples acquired across 11 plates, together with the inclusion of multiple pooled reference sample types, process controls, and system suitability injections, makes it particularly valuable for the development and benchmarking of normalization and batch-correction methods. Because the dataset contains known technical structure across plates in addition to biologically meaningful variation associated with dose, dose rate, and time after exposure, it provides an opportunity to evaluate how effectively different normalization approaches preserve biological signal while removing unwanted technical variation. The availability of both raw and normalized data, together with extensive metadata and reference samples distributed throughout the experiment, also allows testing of new approaches for signal extraction, missing value handling, longitudinal drift correction, and cross-batch harmonization.

## Data Availability

All data presented in the current work are available on Panorama Public^38^ (https://panoramaweb.org/teirex-2a-ldxr-mouse-pelt.url) and were assigned the ProteomeXchange^39^ ID: PXD078423 (https://doi.org/10.6069/5ey2-j220).

## Code availability

MSConvert^18^ is available from https://proteowizard.sourceforge.io/. DIA-NN^21^ is available from https://github.com/vdemichev/DiaNN. Carafe^19,20^ is available from: https://github.com/Noble-Lab/Carafe. Skyline^22,23^ is available from https://skyline.ms/skyline.url.

The single Nextflow workflow used in the current work to search and quantify the MS data presented here is publicly available at: https://nf-teirex-dia.readthedocs.io/en/latest/. This workflow was used to generate empirical spectral libraries using Carafe and DIA-NN, followed by MS data searching using DIA-NN. The workflow also implements a command line version of Skyline to automatically align retention times and impute missing boundaries before performing data extraction and quantification. This same pipeline can also perform precursor and protein level normalization. The precursor data presented here was normalized by this workflow while protein level normalization and batch correction were performed by in house code.

All in house code written for the analysis of the data presented in this work is available online at https://github.com/uw-maccosslab/manuscript-teirex-2a-ldxr-mouse-pelt.

## Author Contributions

M.J.M., C.C.W., P.A.R., D.C., S.E.C. and A.M.S. conceived the experiments. M.J.M., C.C.W. A.Z., G.E.M. A.M.S, J.L.I., and J.H.M. designed experiments. C.C.W., D.P.J.E., A.M., B.A.S., T.N.S., A.M.S, J.L.I., K.H.W., and J.H.M. designed the mouse irradiation protocols and performed mouse work including initial tissue processing. K.N., L.O.-H., J.d.C. and A.M.S. designed and performed X-ray calibration and harmonization experiments at LBNL and C.C.W., N.C., and E.C.F. designed and performed X-ray calibration and harmonization experiments at UW. A.Z. C.C.W., and B.M. performed proteomic sample preparation. A.Z. and G.E.M. performed MS data acquisition. M.R. performed the analysis of proteomic data with help from A.Z., G.E.M, P.A.R. and M.J.M. M.R and N.S. contributed new analytical tools to perform proteomics data analysis. The manuscript was written by A.Z. and M.R. with contributions from all authors. All authors discussed the results and commented on the manuscript. All authors have given approval to the final version of the manuscript.

## Acknowledgements

We would like to thank Vagisha Sharma for help with preparing the Panorama Public data repository for this work. The work presented here was supported by the National Institutes of Health, National Institute of General Medical Sciences under Award P41GM103533 (to M.J.M.) and award 1R21AI194031-01 to J.I. This research is based upon work supported in part by the Office of the Director of National Intelligence (ODNI), Intelligence Advanced Research Projects Activity (IARPA) under Targeted Evaluation of Ionizing Radiation EXposure (TEI-REX) program contract 20008-D2021-2107310008 (LBNL) and through the Army Research Office contract W911NF2220059. The views and conclusions contained should not be interpreted as necessarily representing the official policies, either expressed or implied, of ODNI, IARPA, ARO, or the U.S. Government. The U.S. Government is authorized to reproduce and distribute preprints for governmental purposes notwithstanding any copyright annotation therein.

